# SpacGPA: annotating spatial transcriptomes through *de novo* interpretable gene programs

**DOI:** 10.1101/2025.10.01.679918

**Authors:** Yupu Xu, Lirui Chen, Shisong Ma

## Abstract

Spatial transcriptomes, especially those from high-density platforms (e.g., Visium HD, Stereo-seq, Xenium Prime 5K), enable detailed tissue mapping but remain challenging to annotate for spatial domains and spatially variable genes (SVGs). Most existing methods depend on companion single-cell datasets and are sensitive to cell segmentation. We present SpacGPA, a segmentation-free, single-cell-independent framework that identifies *de novo* interpretable gene programs—co-expressed gene modules capturing key biological processes—from spatial transcriptomes. Leveraging GPU-acceleration and block-matrix operations, SpacGPA efficiently detects gene programs based on a graphical Gaussian model in ultra-large datasets. These programs exhibit domain-specific spatial expression patterns and cell type-specific expression in independent single-cell datasets, serving as robust markers for spatial domains. Applied to four mouse and human spatial datasets, SpacGPA identified 58-109 gene programs per dataset, revealing refined tissue structures. Hub genes within these programs correlate with canonical cell type markers and SVGs, enabling systematic SVG identification. The programs are also functionally enriched and transferable across datasets and platforms. Built on AnnData, SpacGPA provides a scalable, low-hardware solution for high-resolution annotation of tissue organization and gene regulation.

## Background

Spatial transcriptomics (ST) technologies enable high-throughput transcriptome profiling while preserving the spatial organization of tissues, providing powerful opportunities to dissect cellular composition and molecular mechanisms^1,2^. ST analyses typically aim to identify spatial domains, detect spatially variable genes (SVGs), and align multiple sections for three-dimensional reconstruction of tissues or organs^3-5^. Most existing methods, such as RCTD, Tangram, cell2location, rely on companion single-cell datasets for accurate cell type deconvolution and spatial domain annotation^6-8^. Recently, high-resolution ST platforms such as Visium HD, Stereo-seq, and Xenium Prime 5K have advanced spatial resolution to near single-cell or even subcellular levels, enabling transcriptome-wide measurements for hundreds of thousands of cells within a single section^9-11^. This increased resolution allows detailed investigation of the cellular microenvironment and subcellular localization of gene expression, but the resulting data—ultra-high throughput, large scale, and strongly spatially dependent—pose new analytical challenges. Current approaches often require accurate cell segmentation to reconstruct single-cell expression matrices for downstream analysis^12,13^. Despite progress with methods such as Cellpose, Bering, Baysor and SCS, segmentation remains imperfect, underscoring the need for alternative segmentation-free strategies^12,14-16^.

SVGs exhibit marked spatial heterogeneity within tissues and often map to specific anatomical structures or localized functions^3^. Early approaches relied on global spatial autocorrelation metrics, such as Moran’s I, to identify SVGs by quantifying the association between gene expression and spatial proximity^17^. However, these global metrics struggle to disentangle influences of cell-type composition and tissue hierarchy, and they miss genes expressed across multiple discrete regions^18^. Methods such as SpatialDE and SPARK incorporate multi-scale spatial information and improve SVG detection, yet they remain sensitive to data resolution and often lack the sensitivity to detect subtle differences in high-resolution, single-cell–level data^3,4,19^. Moreover, while several approaches incorporate spatial domains, the prevailing gene-by-gene paradigm provides limited statistical leverage to define domain-specific gene sets or to consolidate signals into program-level readouts aligned with tissue architecture^20,21^. This limitation results from neglecting gene–gene interactions and co-expression relationships during SVG identification^19^.

Coupling spatial structure with molecular function and connecting it to other lines of biological evidence is another key requirement in ST analysis^2,22^. For example, gsMap integrates high-resolution ST data with spatial domain annotations and genome-wide association study (GWAS) statistics to spatially localize the distribution of cells associated with complex traits^23^. This approach has revealed differences in the spatial distributions of neurons linked to distinct psychiatric disorders in mouse brain sections and identified pathway genes upregulated in these cells^23^. Nonetheless, a unified framework is still missing. In a single pass, such a framework should define spatial structures, report gene-level signals within each structure, translate them into pathway-level interpretations aligned with tissue architecture, and produce gene-centered outputs to guide validation.

As ST is applied across multiple samples and tissue regions, integrating and transferring information across datasets and platforms have become a major challenge^24^. Slice-alignment methods, such as PASTE, STalign and SLAT, register sections using expression and/or morphology to support cross-slice integration and three-dimensional reconstruction^5,25,26^. Yet cross-platform transfer of spatial patterns is impeded by differences in spatial resolution, capture chemistry, and sequencing depth, so signals are not directly portable^27^. Deconvolving spatial data using single-cell transcriptomes achieves cross-dataset information transfer to some extent (for example, by providing the cellular composition at each location), but these methods are confined to the cell-type level and cannot convey higher-order tissue structure information^6,8^. In addition, how to feed spatial domain information obtained from spatial transcriptomics back into single-cell transcriptomic data —thereby linking dissociated cells with their in situ spatial context as well as provide better annotation of spatial domains—remains an unresolved problem^24^.

To address these challenges, we developed an integrated, easily-accessible, and scalable spatial transcriptomics analysis framework named Spatial and single-cell Gene Program Analysis (SpacGPA). SpacGPA builds gene co-expression networks directly from large-scale spatial data to identify gene programs, highlights hub genes with distinct spatial patterns, and performs functional enrichment analysis to assign biological meaning to each program. The per-spot activity of these programs delineates spatial domains and supports sensitive detection of SVGs within tissue architecture. A gene-centric joint analysis module integrates multiple sections across platforms and resolutions and transfers learned programs to independent spatial datasets and single-cell profiles. SpacGPA runs under limited computational resources, scales to more than one million spots, and integrates with mainstream workflows. The software is freely available at https://github.com/MaShisongLab/SpacGPA, and across diverse datasets, it consistently identifies gene programs and performs downstream analyses, often outperforming existing tools.

## Results

### An Overview of SpacGPA

We developed SpacGPA, a framework to efficiently annotate spatial transcriptomes through *de novo* gene programs. The framework consists of two main components: (i) a GPU-based algorithm for constructing gene co-expression networks and identifying modules using a modified Markov clustering algorithm, and (ii) a suite of tools to annotate these programs and apply them to spatial transcriptomic data **(Figure 1)**.

**Figure 1:**
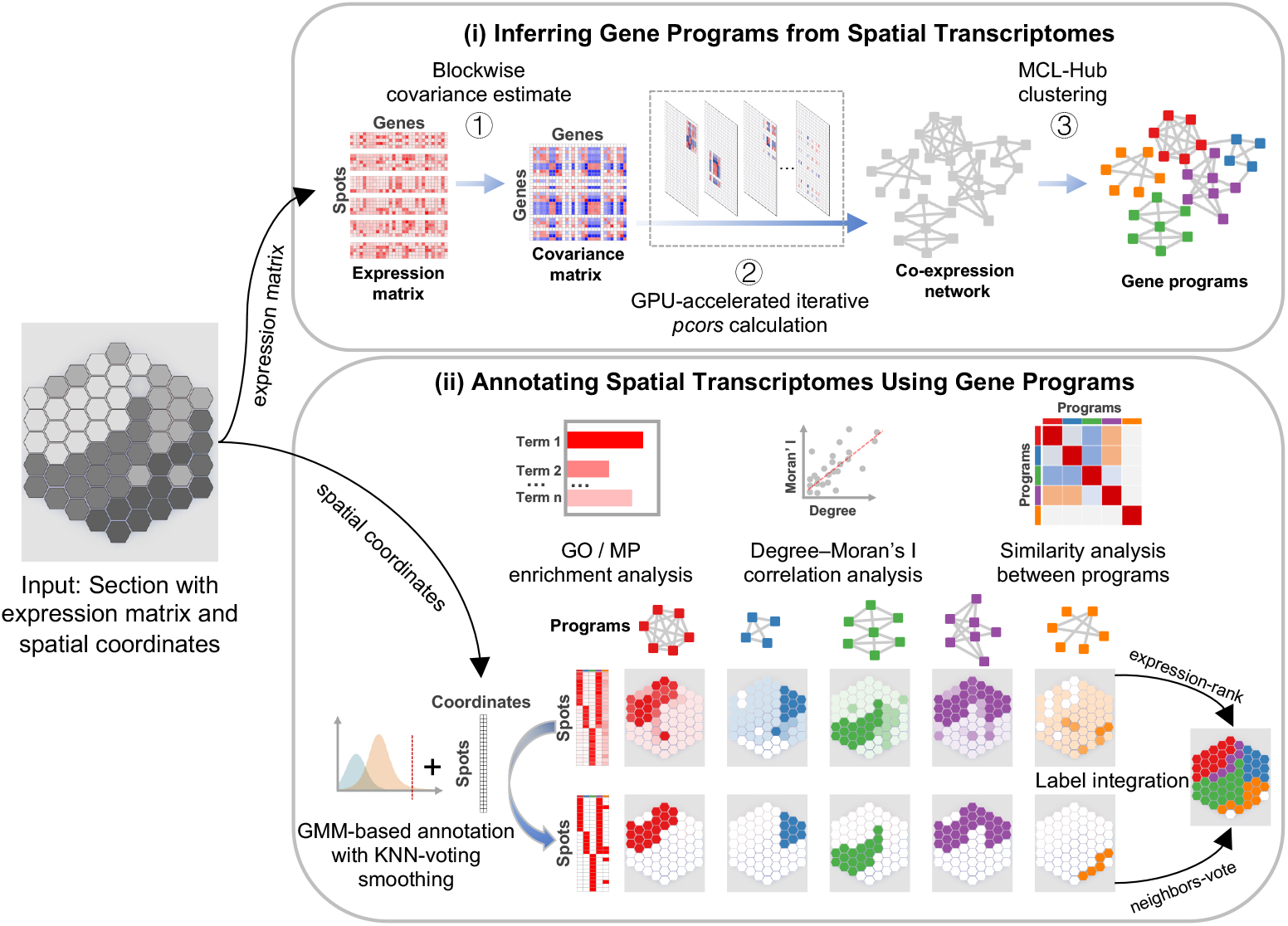
Overview of the SpacGPA workflow. SpacGPA analyzes spatial transcriptomics dataset containing an expression matrix and spatial coordinates. (i) It constructs gene co-expression networks from expression matrix and then identifies gene programs. (ii) It performs functional analysis and defines these programs, scores their expression intensity, and uses them to annotate spots and identify spatial domains.

For network construction, we implemented a Python-based algorithm that calculates gene– gene partial correlations from spatial transcriptomic expression matrices (**Figure 1**). Our previous MATLAB tool, SingleCellGGM, used an iterative, random gene-sampling strategy to estimate co-expression in sparse single-cell datasets but was limited by CPU-based computation, moderate runtimes, and high memory usage for datasets exceeding one million cells^28^. SpacGPA overcomes these limitations with PyTorch-enabled block-matrix operations and by running the same iterative, random gene-sampling strategy on GPUs to compute partial correlations efficiently. Networks of 10k–200k cells can now be constructed in under 30 minutes, and datasets with over one million cells are processed within an hour on a standard server (128 GB RAM, 11 GB GPU). Program detection is performed using MCL-Hub, a GPU-accelerated extension of the Markov clustering algorithm^29^. Both network construction and MCL-Hub require minimal parameter tuning. Unlike many spatial algorithms that use only a few thousand highly variable genes (HVGs), SpacGPA can process nearly all genes (approximately 20,000 after removing low-expression genes detected in fewer than 10 cells)^30^. Although the current focus is on spatial data, this optimized strategy also applies to conventional single-cell datasets, broadening its utility.

The annotation component provides ontology-based functional enrichment (e.g., GO, MP), hub-gene–weighted program expression scoring with visualization, spatial autocorrelation quantification (Moran’s I), and Gaussian-mixture modeling to identify program-positive spots or cells. Additional utilities include automated label assignment, program-level dimensionality reduction for conventional KNN-based clustering, and visualization of global expression patterns and inter-program relationships. Built on the widely used AnnData object, our framework adopts streamlined parameterization and standardized interfaces, ensuring compatibility with mainstream spatial transcriptomics workflows^31,32^. Together, these features enable efficient and interpretable program-based analysis of spatial transcriptomes, including identification of SVGs, partitioning of spatial domains, and cell type annotation, as illustrated in the following examples.

### SpacGPA Identified Refine Structures and SVGs in HD Spatial Data

We first applied SpacGPA to a mouse small intestine Visium HD dataset from 10x Genomics, consisting of 83,337 square spots binned at 16 µm without cell-segmentation. SpacGPA identified 61 gene programs (M1–M61), each representing a distinct gene co-expression module (**Figure 2A, Table S1**). Most programs displayed spatially clustered expression patterns and were enriched for specific biological processes. For example, program M1 (539 genes) was strongly associated with B cell activation (58 genes, *P* = 6.18E-35). Hub genes within M1, defined by intra-module connectivity (degree) (**Figure 2B**), included *Ighd, Cd79a, Cd79b*, and *Cd19*—all classical B cell markers—showing localized expression in structures resembling Peyer’s patches^33,34^. These genes also exhibited high Moran’s I spatial autocorrelation. The program-level expression, calculated as the degree-weighted average of the top 30 genes, closely matched hub-gene expression but showed even stronger spatial autocorrelation, demonstrating that programs can enhance and generalize hub-gene signals, thereby serving as composite spatial markers of cell populations. Spots with positive M1 expression, identified using a Gaussian mixture model (see Methods), were accordingly annotated as B cells (**Figure 2B**).

**Figure 2:**
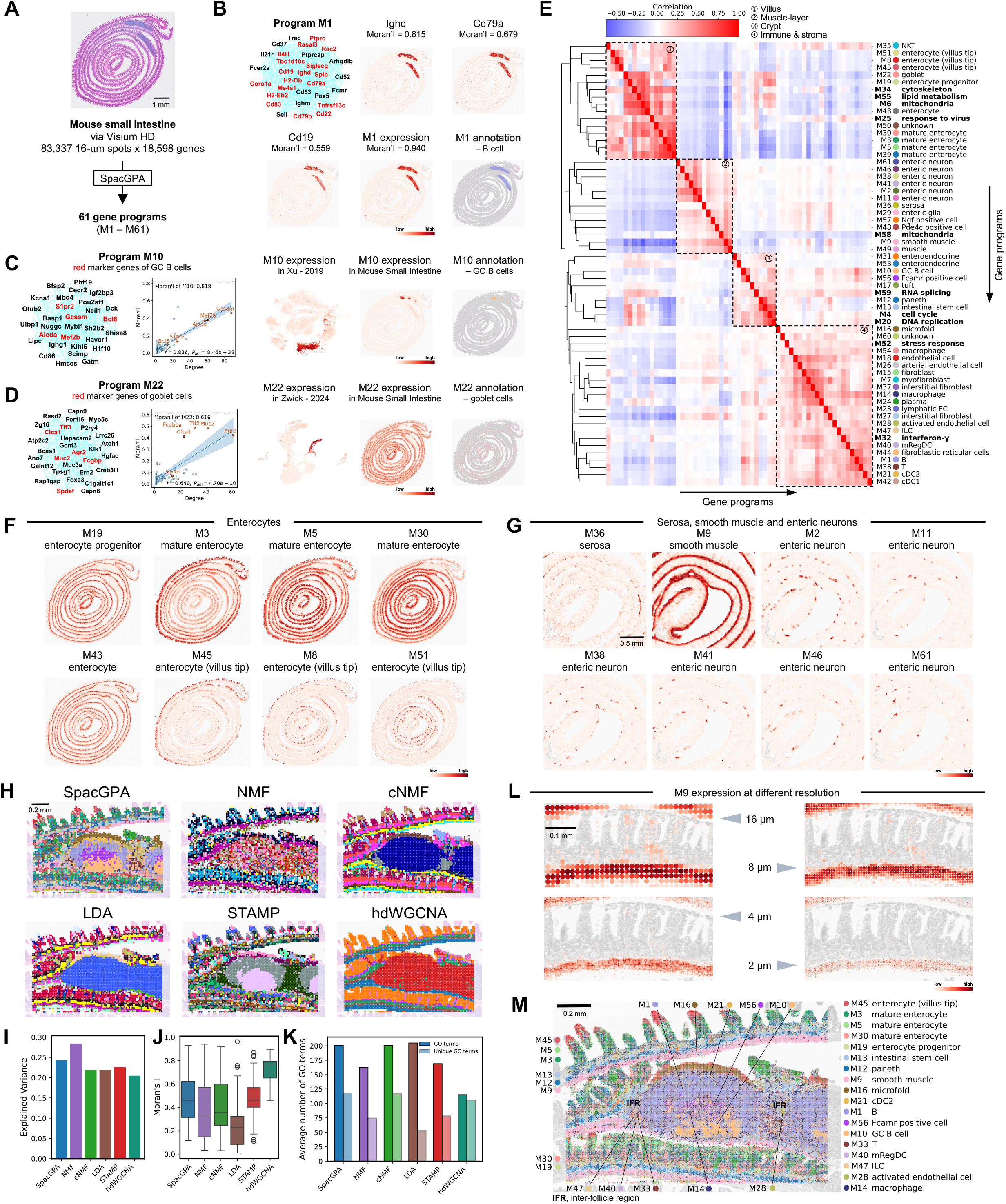
SpacGPA identifies gene programs from mouse small intestine dataset and applies them to SVGs and spatial domains analysis. **(A)** SpacGPA identified 61 programs from the mouse small intestine. **(B)** Subnetwork and spatial expression of program M1 and its genes. Only the top 30 genes with the highest connectivity are shown in the subnetwork due to layout constraints. Spots with high expression of M1 are annotated as B cells. **(C)** Identification of SVGs associated with GC B cells from program M10 and delineation of the corresponding spatial domains. The scatter plot shows the relationship between degree and Moran’s I of genes within M10; regression fitting and 95% confidence intervals are included; r represents the Pearson correlation coefficient; *P* values are corrected by Benjamini-Hochberg; Moran’s I of the program is also reported. Expression distribution of M10 was further validated in a reference single-cell dataset^37^. **(D)** Identification of SVGs associated with goblet cells from program M22 and delineation of the corresponding spatial domains. **(E)** Heatmap showing correlation between the programs. The color bar reflects correlation strength. Labels on the right indicate the cell type or the biological activity assigned to each program; shared activity programs are labeled in bold. **(F)** Spatial expression of programs associated with different intestinal epithelial cells, ordered by differentiation state and spatial location. **(G)** Spatial expression of programs associated with serosal cells, smooth muscle cells, and enteric neurons. **(H)** Integrated annotation integration of the same mouse small intestine dataset using different methods, with a Peyer’s patch region highlighted. Except for SpacGPA, all other methods used the binarization strategy provided by STAMP for annotation integration. **(I)** Total explained variance of programs identified by different methods. **(J)** Distribution of Moran’s I for programs identified by different methods. **(K)** Average number of enriched and deduplicated GO terms for programs identified by different methods. **(L)** Differences in spatial expression distribution of program M9 at multiple resolutions. Within each panel, spot radius scales linearly with program’ expression intensity; values <20% of the panel’s maximum are proportionally reduced to zero, revealing the background where no expression is present. **(M)** Integrated annotation of the mouse small intestine at 4 µm resolution using all programs except the shared activity programs, with a Peyer’s patch region shown. See Figure S3 for annotation of complete section. Scale bars: A, 1 mm; G, 0.5 mm; H, 0.2 mm; L, 0.1 mm; M, 0.2 mm.

Gene programs can be leveraged to identify spatially variable genes (SVGs), since hub genes often align with SVGs. For example, in program M10 we detected hub genes with both high spatial autocorrelation and high module degrees, including *Gcsam, Mef2b, Aicda, S1pr2*, and *Bcl6* (**Figure 2C**). Across all M10 genes, module degree was positively correlated with Moran’s I (*R* = 0.64, *P* = 4.70 E-10), indicating that hub genes tend to be SVGs. Since these hub genes are known germinal center (GC) B cell markers or regulators of GC B cell function, we designated M10 as a GC B cell program^35,36^. This inference was validated in an independent mouse small intestine single-cell dataset, where M10 signal was enriched in a cluster annotated as GC B cells^37^. Similarly, M22 was annotated as a goblet cell program, containing hub genes *Muc2, Tff3, Agr2, Clca1*, and *Fcgbp*, all established goblet cell markers^38,39^. Spots expressing M22 were thus labeled as goblet cells, and this inference was validated in an independent intestinal epithelial single-cell dataset^40^ (**Figure 2D**). For most genes in M22, degrees were positively correlated with Moran’s I. Indeed, in 57 of 61 programs, we observed similar correlations (*R* > 0.5 and P < 0.05; **Figure S1**), demonstrating that hub genes frequently correspond to SVGs, providing a systematic approach to identify spatially variable structures, even though absolute Moran’s I values may vary across programs.

To systematically examine how the 61 gene programs delineate cellular structures in the mouse small intestine, we performed hierarchical clustering based on inter-program expression similarity (**Figure 2E**). Four major clusters emerged: (i) a villus group of 16 programs, including 9 marking distinct enterocyte subgroups; (ii) a muscle-layer group of 13 programs aligning with the muscle band, encompassing serosa (M36), smooth muscle (M9), and six enteric neuron programs; (iii) a crypt group of 10 programs localized to the stem cell region, including cell cycle (M4), DNA replication (M20), intestinal stem cells (M13), Paneth cells (M12), and enteroendocrine cells (M31, M53); and (iv) an immune and stroma group of 22 programs enriched in immune structures such as Peyer’s patches or broadly distributed across intestinal regions.

Several programs further resolve subtle differences within the same cell type, thereby revealing refined structural states. For example, nine enterocyte programs were identified: M19 marked progenitors, M43 marked young enterocytes, while M3, M5, and M30 corresponded to mature enterocytes with distinct spatial dynamics along the proximal-distal axis of small intestine—M3 enriched proximally, M30 distally, and M5 spanning both ends. Additional programs (M45, M8, M51) localized to villus tips but varied in proximal–distal distribution, reflecting spatial–temporal functional heterogeneity^41^ (**Figure 2F**). Similarly, six enteric neuron programs (M2, M11, M38, M41, M46, M61) were identified, interspersed with smooth muscle cells (labeled by program M9), suggesting a complex neuronal network coordinating smooth muscle and intestinal motility^42^ (**Figure 2G**). Together, these results demonstrate that SpacGPA can resolve fine-grained structures within the mouse intestine.

We then used 51 of the 61 programs to annotate the mouse small intestine sample at 16 µm resolution. The remaining programs were omitted because they represent generic cellular activities with limited cell identity information, such as M4 for cell cycle regulation, M20 for DNA replication, and M32 for interferon-γ response. Spots positive for each program were identified using a Gaussian mixture model (GMM). Since a spot could be marked by multiple programs, we developed a scheme to assign each spot a unique final label, prioritizing programs with fewer positive spots (see Methods). This strategy enabled us to generate a comprehensive annotation of the intestine at 16 µm (**Figure S2**).

We next compared SpacGPA with other commonly used program/module-based approaches, including NMF, cNMF, LDA, STAMP, and hdWGCNA^43-47^ (**Figure S2**). For fairness, NMF, LDA, and STAMP were set to infer 61 programs/topics, cNMF identified 40 programs as suggested by its internal criteria, and hdWGCNA produced 13 modules after parameter optimization. Overall, SpacGPA achieved the clearest structural annotation. In a magnified Peyer’s patch region (**Figure 2H**), four of the six methods—but not NMF or hdWGCNA—detected the microfold cell domain, while only SpacGPA and STAMP identified the GC B cell region. SpacGPA further delineated a domain (M56) enriched for *Fcamr*, potentially corresponding to follicular dendritic cells (FDCs)^48^, and resolved multiple immune cell types in inter-follicle regions, including activated ECs (M28), T cells (M33), ILCs (M47), and mRegDCs (M40), as well as cDC2s (M21) in the dome region— features missed by other methods. Performance was quantified using explained variance (EV), where SpacGPA ranked second, with an EV approximately 0.25 (**Figure 2I**). Moran’s I distributions showed similar medians for SpacGPA and STAMP, both only trailing hdWGCNA (**Figure 2J**); however, because hdWGCNA produced far fewer modules, its Moran’s I may be inflated. When comparing the biological interpretability, programs identified by SpacGPA and cNMF were enriched with more unique GO terms than those by other methods (**Figure 2K**).Taken together, SpacGPA achieved more balanced performance across all criteria.

Since Visium HD data can be analyzed at different resolutions, we further tested whether programs from low-resolution analysis (16 µm) could be applied to high-resolution data (2–8 µm). Indeed, program M9 for smooth muscle consistently highlighted muscle-layer structures at all resolutions, but the 4 and 2 µm data aligned more closely with individual cells (**Figure 2L**). We therefore annotated the sample at 4 µm using 51 identity programs, as too many 2 µm spots lacked sufficient counts for annotation (**Figure S3**). At this resolution, magnified views of Peyer’s patches revealed finer details and clearer layer structures than those annotated at 16 µm (**Figure 2M**). Thus, SpacGPA offeres the advantage of enabling direct transfer of gene programs across resolution levels.

### SpacGPA Identifies Functionally Enriched Gene Programs

Beyond spatial domains, ST analyses also aim to identify key genes underlying domain-specific functions and to determine how these genes contribute to corresponding phenotypes^2^. The gene programs identified by SpacGPA bridge spatial structures and gene functions, effectively linking tissue architecture to molecular phenotype. To demonstrate its broad applicability, we applied SpacGPA to an E16.5 mouse embryo dataset generated on the Stereo-seq platform by Chen et al., which contains 121,480 spots binned at 50 µm without cell segmentation^10^. SpacGPA identified 106 distinct gene programs (M1–M106) (**Figures 3A and Table S2**), which delineated key organ and tissue structures. For example, three programs—corresponding to skeletal muscle (M1), smooth muscle (M34), and cardiac muscle (M39)—outlined muscle distribution across the embryo (**Figure 3B**). These programs contained canonical marker genes, including *Neb, Ttn*, and *Acta1* for skeletal muscle; *Myh11, Cnn1*, and *Acta2* for smooth muscle; and *Myh6, Tnni3*, and *Myl7* for cardiac muscle^49,50^. SpacGPA also identified novel structures missed in the original analysis, such as mesothelium (M89), the myotendinous junction (M37), melanocytes (M82), and Schwann cells (M25) (**Figure 3C**). Moreover, the pipeline resolved three distinct meningeal layers surrounding the brain—arachnoid (M5), endothelium (M19 and M54), and pia mater (M55)— compared to a single layer in the original annotation (**Figure 3D**). Notably, two programs marked endothelium: M19 as a general marker for all endothelial cells, and M54 as brain-specific, containing genes critical for blood-brain barrier function, such as *Cldn5, Mfsd2a, Abcb1a, Slc7a5*, and *Agrn*^51,52^ Additionally, as in the previous example, the genes within each of these programs showed a strong correlation between Moran’s I and node degree, supporting the observation that hub genes tend to be SVGs (**Figure S4**).

**Figure 3:**
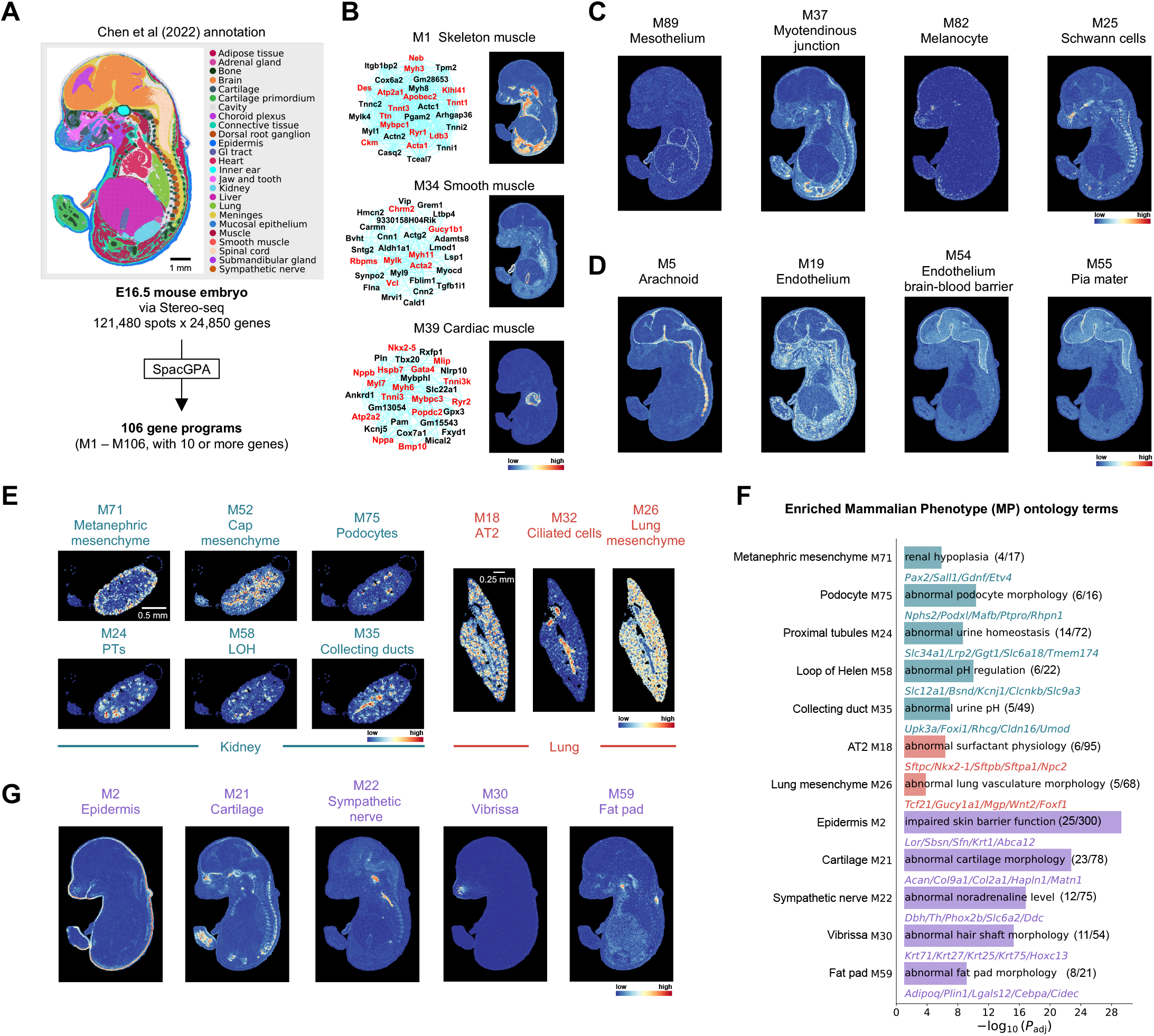
SpacGPA identifies functionally enriched gene programs from a whole-mice embryonic dataset. The dataset, referred to as MOSTA, was obtained from Chen et al.^10^. **(A)** SpacGPA identifies 106 distinct programs from an E16.5 mouse embyo dataset from MOSTA^10^. Spots are colored according to annotations from the original studies. **(B)** Subnetworks and spatial expression of the skeletal muscle program M1, smooth muscle program M34, and cardiac muscle program M39. Corresponding muscle marker genes are highlighted in the subnetworks. **(C)** Spatial expression of the mesothelium program M89, myotendinous junction program M37, melanocyte program M82, and Schwann cells program M25. **(D)** Spatial expression of the arachnoid program M5, general endothelial cell program M19, blood-brain barrier program M54, and pia mater program M55. **(E)** Functionally enriched or spatially specific programs in the kidney and lung identified from MOSTA. **(F)** Representative MP terms enriched by different programs. The number of genes enriched in the corresponding MP and total number of genes in the program are indicated. Genes (partial) with the MP term in the program are listed, ranked by their connectivity degree. *P* values are corrected by Benjamini-Hochberg. **(G)** Spatial expression of the epidermis program M2, cartilage program M21, sympathetic nerve program M22, vibrissa program M30, and fat pad program M59. Scale bars: A, 1 mm; E, left (kidney), 0.5 mm; E, right (lung), 0.25 mm.

SpacGPA also identified programs enriched with genes whose mutations in mouse models lead to phenotypes consistent with the function of the corresponding organ or cell type. In the kidney, six programs were revealed for distinct cell types, including metanephric mesenchyme (M71), cap mesenchyme (M52), podocytes (M75), proximal tubules (M24), the loop of Henle (M58), and collecting ducts (M35) (**Figure 3E**). Based on Mammalian Phenotype (MP) ontology enrichment analysis, five of the six programs were significantly enriched for relevant phenotypes^53^. For instance, 14 of 72 genes in M24 have been shown to cause abnormal urine homeostasis when mutated (*P* = 2.03E-08), including *Slc22a12, Slc34a1*, and *Lrp2*, consistent with proximal tubule function^54,55^; and 6 of 16 genes in M75 cause abnormal podocyte morphology when mutated (*P* = 3.99E-10), including *Nphs2, Podx*, and *Rhpn1*, confirming podocyte specificity^56,57^ (**Figure 3F**; see **Table S2** for a complete list). Similarly, among eight programs for lungs and other structures, all but one (M32) were enriched for MP terms corresponding to phenotypes observed in mutants, including impaired skin barrier, abnormal cartilage morphology, and altered noradrenaline levels (**Figure 3E–G**).

We further applied SpacGPA to an ultra-large mouse pup dataset measured on the Xenium Prime 5K platform, containing 917,936 segmented cells. Unlike sequencing-based platforms, Xenium Prime 5K uses imaging to measure ∼5,000 genes. SpacGPA identified 88 gene programs (M1–M88), capturing major and refined tissue structures (**Figures 4A, S5; Table S3**). For example, six programs outlined the layered structures of the eye—including lens, ganglion cell layer, retinal neuronal layer, photoreceptor layer, pigment cell layer, and choroid (M58, M80, M7, M37, M60, and M51)—while two programs corresponded to ear structures (M61, M62), and seven programs marked cartilage and bone-related structures (M63, M71, M53, M48, M49, M68, M79) (**Figure 4B**). Many programs were enriched for relevant MP terms; for instance, 12 of 19 genes in M61 have been shown to cause impaired hearing when mutated (e.g., *Otof, Pou4f3, Slc17a8*^58^), while M62, expressed in the membranous labyrinth containing otoliths critical for balance, includes 4 genes (*Nox3, Noxo1, Oc90, Slc26a4*^59,60^) whose mutation decreases otolith number, suggesting additional candidate genes for the same phenotype (**Figure 4C**).

**Figure 4:**
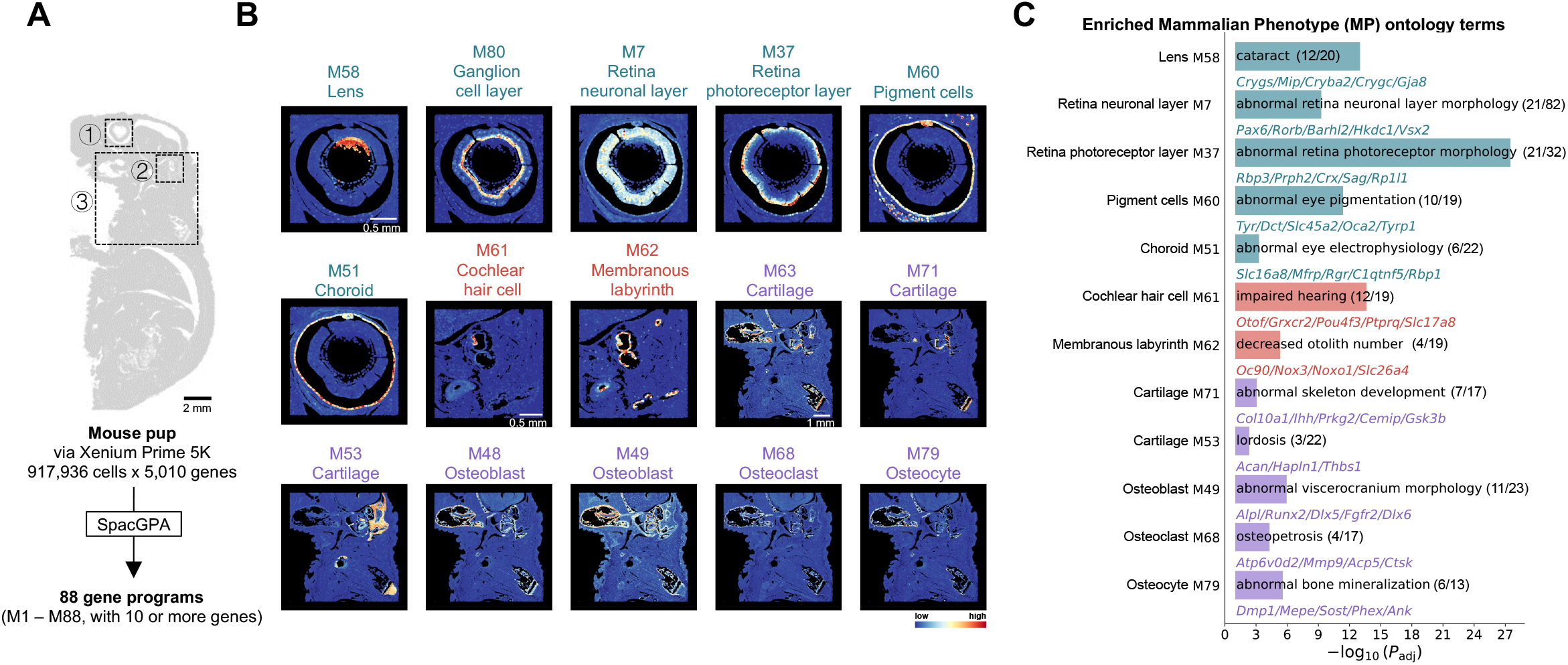
SpacGPA identifies spatially continuous programs from a mouse pup section. **(A)** SpacGPA identified 88 programs from the mouse pup dataset, a whole-body newborn mouse sagittal section profiled with 10x Genomics Xenium Prime 5K at single-cell resolution. Three distinct regions are marked:(1) the eye region, (2) the inner ear region, and (3) the skeletal development-related region. **(B)** Representative MP terms enriched for vision-related, inner ear programs, and skeletal development-related programs. **(C)** Spatial expression of vision-related, inner ear, and skeletal development-related programs in their respetive regions. Scale bars: A, 2 mm; B (top/middle/bottom; corresponding to regions 1–3 in A), 0.5 mm / 0.5 mm / 1 mm.

Overall, SpacGPA identified gene programs for both general and refined structures in mouse embryo and pup spatial transcriptomes derived from different spatial technology platforms. These programs are functionally enriched, capturing both known markers and novel candidate genes for future investigation.

### SpacGPA Enables Multi-Sample Analysis and Cross-Sample Transferability

As spatial transcriptomics becomes more widely applied, joint analysis of multiple samples and knowledge transfer across datasets or platforms become increasingly important^2,27^. SpacGPA supports multi-sample joint analysis, and the resulting gene programs serve as a vehicle to transfer information between datasets. We applied SpacGPA to six mouse brain coronal sections measured with first-generation low-resolution Visium (V1/V2) (**Figure 5A**). Unlike Visium HD, V1/V2 spots have a diameter of ∼55 µm, each potentially containing up to a dozen cells, spaced ∼100 µm apart, yielding 2,000–5,000 spots per slide^61^. Joint analysis combined these data for a total of 16,603 spots, which improved network construction (larger sample sizes >10,000 spots tend to enhance network inference).

**Figure 5:**
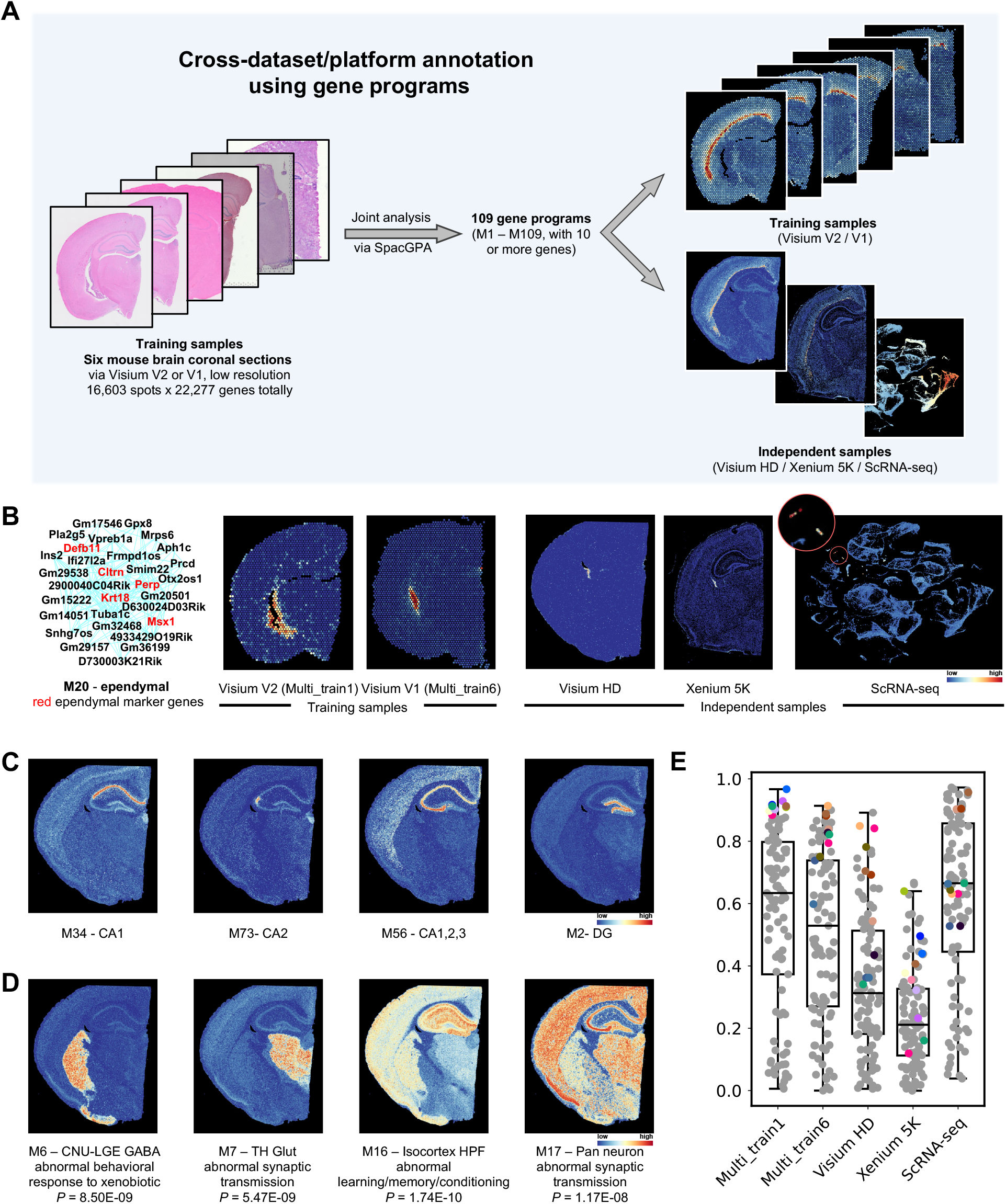
SpacGPA identifies programs from multiple mouse brain sections and transfers them to independent datasets. **(A)** SpacGPA merged six Visium low-resolution coronal sections of the adult mouse brain from 10x Genomics, identifying 109 gene programs from a total of 16,603 spots and 22,277 genes. The spatial expression of program M25 is shown in the training sections and in independent datasets. **(B)** Subnetwork of the ependymal program M20, along with its spatial expression in two training slices and in three independent high-resolution datasets. **(C)** Spatial expression of hippocampal CA1 program M34, CA2 program M73, CA1–CA3 pyramidal cell layer program M56, and dentate gyrus granule cells program M2 in a Visium HD mouse brain section. **(D)** Spatial expression of the GABAergic inhibitory circuit-related program M6, tglutamatergic synaptic transmission program M7, learning/memory/conditioning-related program M16, and whole-brain synaptic activity program M17 in a Visium HD mouse brain section. **(E)** Distribution of Moran’s I in two training slices and in three independent high-resolution datasets for programs identified via joint analysis. Color-highlighted points represent the top 10 programs with the highest Moran’s I values in Multi_train1.

The joint analysis identified 109 gene programs, capturing similar structures across the six training samples (**Figure S6 and Table S4**). These programs also transferred effectively to other platforms of higher resolution (Visium HD, Xenium 5K) and even to single-cell datasets. For example, program M20, enriched with ependymal markers such as *Krt18* and *Msx1*, localized to narrow band-like structures in the lateral and third ventricles, corresponding to the choroid plexus^62,63^ (**Figure 5B**). In independent Visium HD and Xenium 5K samples, M20 highlighted the same structure, and in a single-cell dataset it remained specifically expressed in ependymal cells^64^. This demonstrates that cell type–specific marker programs can be identified even from spots containing multiple cells. Given this robust transferability, we leveraged a deeply annotated mouse whole-brain single-cell atlas to guide functional annotation of the programs^65^.

Other programs similarly highlighted defined brain regions (**Figure S7**). In Visium HD samples, M34 marked the hippocampal CA1; M73 marked CA2; M56 spanned the CA1–CA3 pyramidal cell layer; and M2 highlighted dentate gyrus granule cells (**Figure 5C**). M6 showed high expression at the caudate–globus pallidus border, corresponding to basal ganglia GABAergic circuits, and is enriched for the MP term “abnormal behavioral response to xenobiotic”^66^. M7 was restricted to thalamic nuclei and enriched for the MP term “abnormal synaptic transmission”, reflecting its role in sensory relay^67^ (**Figure 5D**). Programs spanning multiple regions were also identified: M16 covered the cerebral cortex, striatum, and hippocampus, associated with “abnormal learning/memory/conditioning”, while M17 was broadly expressed across neuronal layers, also enriched for “abnormal synaptic transmission”, indicating involvement in brain-wide synaptic activity and signal integration^68^.

Overall, the programs showed consistent spatial localization, as quantified by program-level Moran’s I. Highly variable programs in low-resolution V1/V2 slides also displayed high Moran’s I in scRNA-seq (Moran’s I calculated according to UMAP coordinates), Visium HD, and Xenium 5K (**Figure 5E**). Interestingly, Moran’s I generally decreased as resolution increased, likely reflecting finer spatial details revealed by high-density platforms^30^.

### SpacGPA Enables Refined Analysis of Tumor Samples

Because SpacGPA is built upon the AnnData format, it compatible with many mainstream spatial transcriptomics workflows^31,32,69^. To demonstrate its utility, we applied SpacGPA, along with other tools, to a human lung adenocarcinoma sample measured using the 10x Genomics Xenium Prime 5K platform, comprising 217,922 segmented cells. SpacGPA identified 58 gene programs, for which a deeply annotated human lung single-cell dataset was used to guide functional annotation^70^ (**Figures 6A, S8 and Table S5**). For each program, expression was quantified to construct a “cell-by-program” feature matrix, which was subsequently normalized and scaled to generate a low-dimensional representation of cells. Leiden clustering based on this representation partitioned the cells into 45 clusters, revealing distinct spatial domains (**Figure 6B**). InferCNV analysis indicated that five clusters (2, 5, 9, 31, 41) exhibited elevated copy number variation (CNV) scores relative to other clusters, coinciding with high expression of canonical lung adenocarcinoma epithelial markers (*NKX2-1, EPCAM, MUC1, EGFR*)^71^. These clusters were therefore designated as malignant tumor cells (**Figure 6C**). Comparison of gene program activity across these malignant clusters revealed subgroup-specific signatures. For instance, program M10 was enriched in DNA replication genes; M58 was characterized by coordinated metabolic rewiring and RNA processing; M48 marked an ER-proteostasis/UPR program. These findings indicate that distinct tumor subgroups are transcriptionally specialized, representing multiple differentiation states within the same tumor^72,73^ (**Figure 6D**).

**Figure 6:**
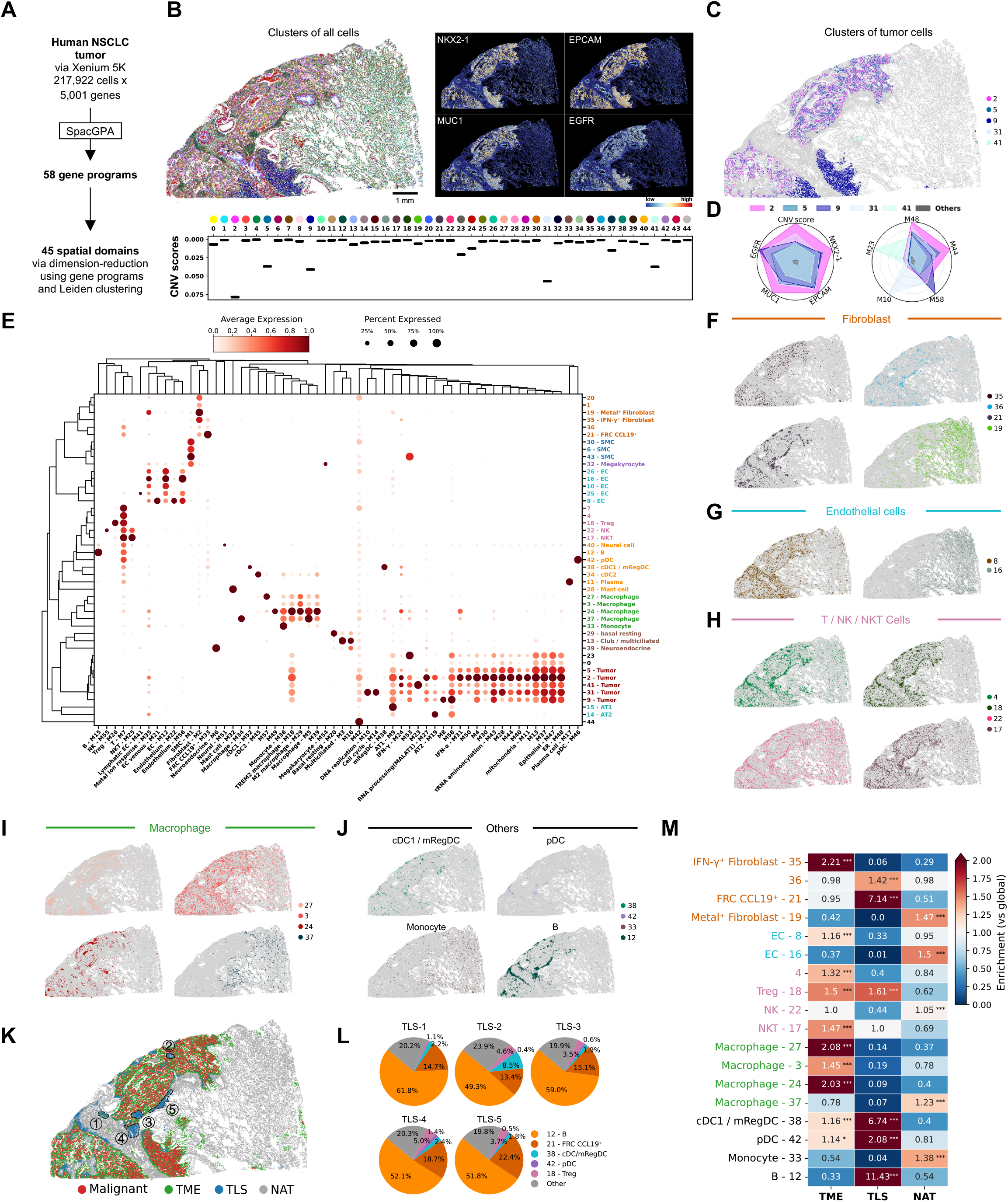
SpacGPA performs refined analysis in human lung adenocarcinoma tissue section alongside other tools. **(A)** SpacGPA identified 58 programs in NSCLC, a human lung adenocarcinoma section with approximately 278,000 spatially resolved cells profiled by 10x Genomics Xenium Prime 5K. Leiden clustering based on the program’s normalized feature matrix divided all cells into 45 spatial domains. **(B)** Identification of tumor cell clusters based on the expression of malignant cell marker genes and the copy number variation scores within each cluster. **(C)** Spatial distribution of tumor cell clusters. **(D)** Normalized mean expression of malignant cell marker genes (left) and tumor cell-specifically programs (right) across different tumor cell clusters. **(E)** Differential expression of programs within each cluster. Bubble size indicates the proportion of cells in that cluster expressing that program, as derived from a GMM model (see Methods); the color bar corresponds to the average expression of the program within cluster. Clusters are annotated by the programs they expressed. Hierarchical clustering of the clusters and programs wer performed based on expression similarity. **(F-J)** Spatial distribution of four fibroblast, two endothelial cells, four T cells, four macrophages, and some other immune cells clusters in tissue section. **(K)** Cell distribution across tissue sections, categorized as malignant cells (Malignant), tumor microenvironment (TME), tertiary lymphoid structures (TLS), and normal adjacent tissue (NAT). Five TLSs are circled according to the number of cells they contain. **(L)** Main cell types and their proportions within five TLSs. **(M)** Heatmap showing fold enrichment of cell clusters in different regions. Color bars reflect enrichment; numbers on squares indicated fold enrichment. Results were analyzed using Fisher’s exact test with Benjamini-Hochberg correction (^*^, *P* < 0.05; ^**^, *P* < 0.01; ^***^, *P* < 0.001). Scale bar: B, 1 mm.

Hierarchical clustering of program expression further organized the cell clusters and programs according to shared expression patterns (**Figure 6E**). Tumor cell clusters formed a discrete branch and were characterized by high expression of programs containing key tumor-associated genes (e.g., *P4HB, CLPTM1L* and *CKAP4* in M48; *MUC1, MUC4*, and *FOLR1* in M13; *IGF2BP3, TUBB3*, and *MAGEC2* in M23)^74,75^. Additionally, tumor cells showed elevated activity of M43, a program enriched for tRNA aminoacylation genes. This observation is consistent with previous reports that tRNA aminoacylation pathways are upregulated in tumors to support the increased translational demand of rapidly proliferating cells^76,77^.

Beyond the malignant population, distinct stromal and immune subsets were also resolved, including fibroblasts, endothelial cells, smooth muscle cells, T cells, NK/NKT cells, and macrophages, and others (**Figure 6E**). Six fibroblast clusters (1, 19, 20, 21, 35, 36) were identified. All of these clusters expressed M2, a general fibroblast program containing markers such as *PDGFRA, POSTN*, and *THBS2*^78^. Additional programs distinguished fibroblast subtypes: M33 in cluster 21 contained canonical fibroblastic reticular cell (FRC) markers (*CCL19, THY1, VCAM1*) and was annotated as FRCs, frequently observed in tertiary lymphoid structures (TLSs)^79,80^; cluster 19 expressed M35, enriched in metal ion response genes; and cluster 19 also expressed M24, an interferon response program. Based on their spatial distributions, clusters 33 and 36 were annotated as cancer-associated fibroblasts (CAFs), cluster 21 as FRCs, and cluster 19 as fibroblasts predominantly located in normal adjacent tissue (NAT) (**Figure 6F**). Other stromal and immune cell populations also displayed heterogeneous spatial patterns across the tumor core, invasive margin, and NAT regions (**Figure 6G–J**).

Integrating information about tumor cell proximity and known TLS marker genes enabled delineation of the tumor microenvironment (TME) and TLS regions. Cells were classified into four categories: malignant tumor cells, TME cells, TLS cells, and NAT cells (**Figure 6K, Methods**). Five prominent TLS aggregates were identified at the tumor border and designated TLS-1 through TLS-5 in order of increasing cell number (Figures 6K, 6L). TLS composition analysis suggested progression from “initial” to “mature” states^81^. B cells predominated in all TLSs; FRCs increased progressively from TLS-1 to TLS-5, reflecting stromal recruitment and matrix remodeling during maturation; dendritic cells peaked in TLS-2, consistent with their early role in antigen presentation; and regulatory T cells were most abundant in TLS-4, in line with the need for immunoregulation in mature germinal centers^82,83^ (**Figure 6L**). Region-specific enrichment analysis further supported these trends: TLSs were significantly enriched in B cells, dendritic cells, and FRCs, whereas different fibroblast subtypes, epithelial cells, and macrophages displayed divergent spatial distributions across tumor core, invasive margin, and NAT (**Figure 6M**).

In summary, this case study establishes a spatial multiscale analysis workflow for lung tumor using SpacGPA. The gene program–based framework enables robust identification of malignant, stromal, and immune subtypes, dissection of TLS maturation, and functional stratification of tumor subgroups, providing a broadly applicable strategy for spatial transcriptomic studies of complex disease tissues.

## Discussion

SpacGPA provides a unified framework that integrates *de novo* gene program discovery with spatial domain identification, distinguishing it from existing tools. Unlike single-gene methods such as SpatialDE and SPARK, which generate lists of spatially variable genes (SVGs) requiring post hoc clustering, SpacGPA directly identifies co-expressed gene programs linked to spatial domains and highlights hub genes^3,4^. Since hub genes generally align with high Moran’s I, we systematically designate those with higher Moran’s I as SVGs, providing a principled way to detect structure-specific SVGs. This approach adapts naturally to structural heterogeneity, as Moran’s I values vary across different domains. Compared with other program-based approaches such as cNMF, STAMP, and LDA—which assign genes fractionally to programs—SpacGPA uniquely assigns genes to single programs, yielding clearer interpretability^44-46^. It also supports functional enrichment analyses, directly linking programs to their biological roles. Benchmarking showed that SpacGPA delivers more balanced performance, uncovering fine structures often missed by other methods (**Figure 2**). Importantly, SpacGPA requires no cell segmentation or reference datasets, runs efficiently on GPUs for million-cell data, and demands minimal parameter tuning, lowering the usage barrier and broadening its utility.

Applications across diverse datasets underscore its robustness and capacity to resolve fine-grained spatial domains. In mouse small intestine, SpacGPA identified 61 programs, including nine enterocyte programs capturing heterogeneity along crypt–villus and proximal–distal axes, enteric neuron networks interwoven with muscle, and diverse immune programs within Peyer’s patches **(Figure 2)**. In embryonic brain, it resolved the multilayered meninges and a blood–brain barrier endothelial program that had been collapsed in prior annotations **(Figure 3)**. In a million-cell mouse pup dataset, it reconstructed fine structures of eye, ear, cartilage, and bone development **(Figure 4)**. Across six adult mouse brain sections, SpacGPA recovered 109 programs representing conserved neuroanatomical regions, which transferred robustly across platforms and resolutions. For example, an ependymal program detected in low-resolution Visium was re-identified in Visium HD, Xenium and scRNA-seq, demonstrating strong reproducibility and efficient cross-platform transfer **(Figure 5)**. Additionally, these programs were enriched for phenotype-relevant genes, underscoring their potential to reveal spatial domain- or structure-specific functions.

SpacGPA also demonstrates translational value. In a human lung adenocarcinoma sample, it resolved tumor core, stroma, and TLSs **(Figure 6)**. Programs enriched in B cells and FRCs localized to tumor margins, corresponding to pathologically defined TLSs and were linked to improved immunotherapy response^81^. By combining clustering with program annotation, SpacGPA highlighted regulatory genes and distinct cell states underlying tumor heterogeneity, enabling refined stratification of the tumor microenvironment. Such capability to map functional modules and structural complexity in situ underscores its potential for biomarker discovery, prognostic assessments, and precision medicine.

In summary, SpacGPA fills a critical gap by unifying functional program discovery with spatial analysis. Its ability to delineate refined structures, identify hub genes as SVGs, avoid segmentation and references, scale efficiently with GPU acceleration, and maintain compatible with mainstream workflows makes it broadly applicable. These features, together with robust cross-platform transferability, establish SpacGPA as a valuable tool for spatial biology and translational research.

## Methods

### SpacGPA Framework

We developed SpacGPA, a comprehensive analysis pipeline for spatial transcriptomics that integrates co-expression network construction, gene program identification, and functional enrichment analysis. SpacGPA builds on the concept of SingleCellGGM (a MATLAB algorithm we previously released) for co-expression network inference based on the graphical Gaussiam model (GGM), but reduces computational burden and accelerates processing through blockwise computation and a GPU-based PyTorch implementation^28^. In addtion, SpacGPA supports the identification of spatially variable genes and delineation of spatial domains based on gene programs.

### Inference of gene co-expression network

To identify gene programs from spatial transcriptomics data, SpacGPA first estimates partial correlations (*Pcor*) between gene pairs to construct a co-expression network. This requires computing both the gene-gene covariance matrix and the co-expression count matrix (i.e., the number of spots co-expressing a given gene pair). For large dataset (>1,000,000 spots), naïve computation impose prohibitive memory costs. To address this, SpacGPA implments a bolckwise strategey that splits matrix operation into menageable chunks processed sequentially^84^.

1. Blockwise covariance estimation Let the expression matrix, after log-normalization and low-expression genes removal, be *X* ∈ ℝ^*n*×*G*^ with *N* spots (observations) and *G* genes (variables). The covariance matrix Σ ∈ ℝ^*G*×*G*^ is defined as:

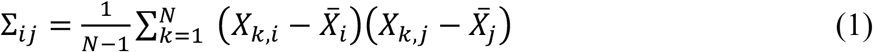

where 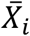 is the mean expression of gene *i*. To reduce memory usage, SpacGPA compustes Σ in blocks. Specially, *X* is split into row blocks *X*^(*p*)^, each containing all *G* genes for a subset of spots. For each block, we compute the partial covariance product:

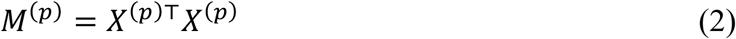

Summing across all blocks yields the product-sum:

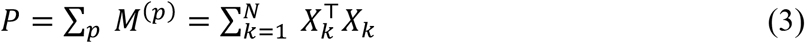

In parallel, we accumulate the row-sum vector *s* and compute its mean 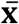:

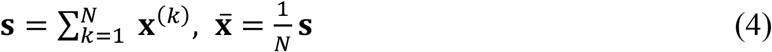

Finally, the covariance matrix is computed as:

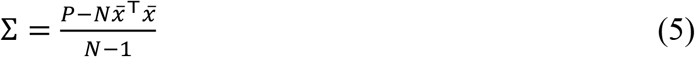

where *P* is obtained from Eq. (3) and 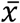 is drived from Eq. (4).
2. Blockwise co-expression count matrix computation SpacGPA also computes a co-expression count matrix *C*, where each entry *C*_ij_ records the number of spots co-expression genes *i* and *j*. First, *X* is binarized to *B* (values >0 set to 1). Then:

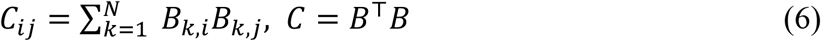

As with covariance estimation, we use a blockwise approach. Each blocks *X*^(*p*)^ is binarized to *B*^(*b*)^, and we compute:

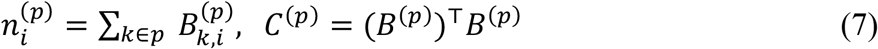

Aggregating across blocks gives:

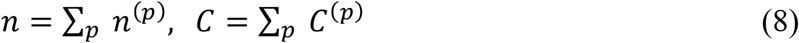

This strategy avoids large intermediate matrices, coupled with GPU-accecelation, enabling eifficient computation even for million-spot datasets.
3. Gene co-expresion network inference SpacGPA infers gene co-expression networks using an iterative random sampling strategy adapted from SingleCellGGM^28^. In each iteration, a subset of *N* genes was sampled (*N* = 2000 if *G* ≥10,000; otherwise, 20% of genes). The sub-covariance matrix for these genes Σ_*s*_ was inverted, and *Pcors* were calculated as:

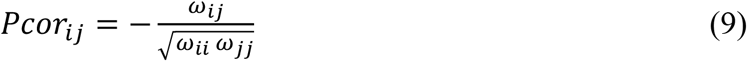

where,

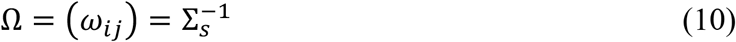

After sufficient iterations, with each pair sampled ∼100 times on average, the lowest absolute *Pcor* for each gene pair across all iterations was taken as the final value. Gene pairs with *Pcor* above a chosen cut-off and co-expression in ≥10 spots were retained for network construction.

To control false discovery, we permute the expression matrix by shuffling values spot-wise within each gene, recalculate *Pcors*, and count gene pairs that simultaneously meet the cut-off and ≥10-spot co-expression. These gene pairs were considered false positives, and FDR was estimated as the ratio of false positives to observed pairs. The final *Pcor* cut-off was chosen as the smallest value achieving an estimated FDR ≤ 0.05, but not below to 0.02. Gene pairs passing these criteria define the co-expression network.

### MCL-Hub algorithm for gene program identification

After constructing the co-expression network, SpacGPA infers gene programs by clustering with the MCL-Hub algorithm, a GPU-enabled extension of the Markov Clustering Algorithm (MCL) with topological refinement^29^. The procedure has three stages:

1. Preprocessing: Build an adjacency matrix *A* from significant gene pairs. Nodes represent genes, edges connect pairs with partial correlation above the threshold and sufficient co-expression, and edge weights equal the partial correlation. Self-loop weights are added on the diagonal to stabilize clustering.
2. Clustering: Apply MCL to matrix *A*, alternating expansion (matrix power) and inflation (element-wise power and normalization) to simulate random walks and amplify strong connections. Iteration continues until convergence, yielding a matrix whose nonzero blocks define candidate modules.
3. Post-processing: Refine modules by addressing singletons and weakly connected genes. The highest-degree node in each module is designated as an attractor. Isolated or zero-degree genes are tested for shortest paths to the attractor, and if supported, merged back into the module. Remaining genes are reassigned by neighbor majority in the original network. Modules with overly simple structures (e.g., stars, chains) are discarded.

### Connectivity-weighted quantification of program expression intensity

To score each program’s expression in each spot, SpacGPA computes a connectivity-weighted sum of its top 30 genes. For a given gene program *p* containing the set of genes *G*_*p*_, we define *G*_*p*30_ as the subset of the top 30 genes ranked by connectivity. The expression intensity *E*_*p*_ of program *p* at spot *s* is defined as:

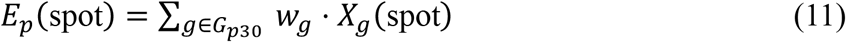

where *X*_*g*_(spot) is the expression level of gene *g* at spot *s*, and *w*_*g*_ is the weight for gene *g* in program *p*. SpacGPA assigns weights based on each gene’s connectivity (degree) within the program’s sub-network, normalized:

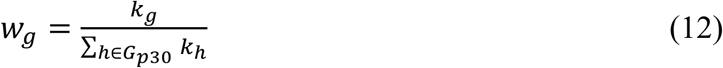

where *k*_*g*_ is the connectivity (degree) of gene *g* in program *p*, and 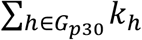 is the sum of connectivities for the top 30 genes in this program.

### Spatial autocorrelation

To assess spatial structure, SpacGPA computes Moran’s I for each gene and program. Moran’s I is computed as:

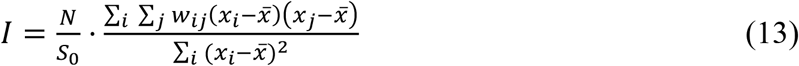

where *x*_*i*_ is the observed value at spot *i* (e.g., gene expression level or program signal intensity), 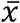 is the mean across all spots, *W*_*ij*_ are elements of the spatial weight matrix *W* (defining neighborhood relationships between spots), *N* is the number of spots, and *S*_0_ = ∑_*i*_ ∑_*j*_ *w*_*ij*_ is the sum of all weights. Moran’s I quantifies the degree of spatial clustering, essentially a weighted correlation between neighboring values normalized by variance. A positive Moran’s I indicates clustering of similar values.

### Dimensionality reduction using program-based features

SpacGPA supports using gene program expression as as a low-dimensional feature space. After scoring each program’s expression in each spot, SpacGPA stores both the raw and normalized program expression matrices in the data object. These can then be used with external tools (e.g., Scanpy^31^) for downstream analyses such as KNN graph constrution, dimensionality reduction (t-SNE^85^, UMAP^86^, etc.), and clustering in the program-based feature space.

### Functional enrichment analysis and annotation of gene programs

Each gene program is functionally annotated through enrichment analysis. SpacGPA performs one-tailed hypergeometric tests to assess over-representation of Gene Ontology (GO) and Mammalian Phenotype (MP) terms. The enrichment p-value is calculated as:

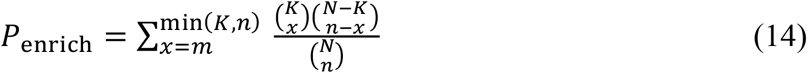

where *N* is the total number of background genes, *K* is the number of background genes associated with the GO or MP term of interest, *n* is the number of genes in the program, and *m* is the number of genes in the program annotated with that term. Programs are annotated by combining enriched GO/MP terms with known marker genes and their spatial pattern. A species-specific GO reference can be built to support custom gene sets.

### Annotating spots using gene programs

Spots are annotated by converting program intensities to binary “active” labels via a two-component Gaussian mixture model (GMM)^87^. For each gene program *p*, we assume its intensity distribution across spots is a mixture of an “inactive” and an “active” Gaussian. We fit a two-component GMM:

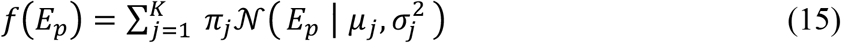

where *π*_*j*_ is the weight of the *j*th Gaussian component (*j* = 1,2), and 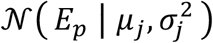 is a Gaussian density with mean *μ*_*j*_ and variance 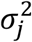. The parameters *π*_*j*_*p μ*_*j*_*p σ*_*j*_ of the mixture are estimated via the Expectation-Maximization algorithm, fitting the GMM to the program’s empirical intensity distribution. For each spot *i*, the GMM yields the posterior probability *P*_*i,h*_ = *P*(*z*_*i*_ = *h*∣*E*_*p*_(*i*)) that the spot belongs to the active component. Spots are labeled as active if *P*_*i,h*_ > 0.0000, ensuringonly those with confidently high program expression are marked. Programs without a clear bimodal distribution do not annotate any spots.

SpacGPA’s GMM-based annotation assigns each spot a binary label (0/1) for each program, but raw labels may contain isolated single-spot assignments or jagged boundaries due to noise. To reduce such artifacts, we apply spatial smoothing. A K-nearest-neighbors graph is built on spot coordinates, and two rules were applied: (i) any spot labeled active spots whose neighbors largely lack that label are switched off, removing spurious isolated spots; (ii) inactive spots surrounded by a majority of active neighbors (e.g. >50%) are switched on, filling “holes” in regions. Neighborhood size and thresholds are platform-dependent (for example, 24 nearest neighbors on Visium). This KNN-voting smoothing produces cleaner, contiguous regions of program activity.

### Label integration

The GMM-based annotation allows each spot to carry multiple program labels (i.e., a spot can be “active” for several programs simultaneously). SpacGPA provides a label integration strategy that combines neighborhood consensus and expression ranking to assign a unique label per spot. First, if exactly one program is active at a spot, that program becomes the spot’s label. If none are active, the spot remains unlabeled. When multiple programs are active, conflicts are resovled using local consensus (Optionally) and expression strength. If local consensus is considered, we calculate the fraction of the spot’s neighbors (in the KNN graph) labeled by each program; if exactly one program dominates (for example, >90% of neighbors) and uniquely leads, it is assigned. If no single program meets this consensus criterion or if local consensus is not considered, we calculate the spot’s expression rank in each candidate program (ranking of the expression intensity among all spots expressing the program, thereby avoiding scale/dispersion bias) and assign the top-ranked program as the unique label. This approach ensures each spot receives a single label in a spatially coherent and quantitative manner.

### Similarity analysis between programs

SpacGPA provides two complementary measures to quantify similarity between program pairs:

1. Expression correlation (Pearson’s *r*). This metric assesses whether two programs share similar spatial expression patterns. The Pearson correlation coefficient *r*_*pq*_ between the spatial intensity vectors of program *p* and program *q* is calculated as:

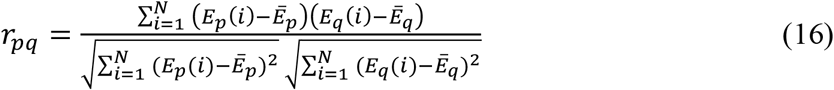
2. Annotation overlap (Jaccard index). This metric compares the sets of spots annotated by each program. For each program *p*, after GMM-based thresholding (and optional smoothing), the set of active spots is *S*_*p*_. Similarly, *S*_*q*_ for program *q*. The Jaccard similarity between programs *p* and *q* is defined as:

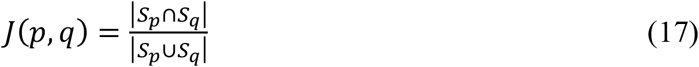

where |*S*_*p*_ ∩ *S*_*q*_ | is the number of spots active for both program *p* and *q*, and |*S*_*p*_ ∪ *S*_*q*_| is the number of spots active for at least one of the two programs. The Jaccard index ranges from 0 to 1.

Together, Pearson correlation and Jaccard overlap provide a comprehensive assessment of the relationships between gene programs in both continuous expression space and discrete spatial domains.

### Joint analysis of multiple sections

SpacGPA supports joint analysis of multiple tissue sections, enabling robust identification of gene programs across samples. Users can select either the intersection or the union of genes across datasets as the gene universe for the analysis. The workflow first extracts the specified gene set (ensuring a consistent gene order across datasets), then aligns each dataset’s expression matrix by these genes and concatenates them along the spot/cell dimension into one combined matrix. Low-expression genes in the merged matrix (for example, expressed in fewer than 10 spots) are filtered out. The resulting matrix is then used for co-expression network inference and program identification as described above. Once gene programs are identified from the joint dataset, all previously described functions —such as computing program expression and annotations —can be applied to individual datasets, allowing comparison of program activity across different sections or experiments.

### Data Access and Processing

We applied SpacGPA to multiple public spatial transcriptomic datasets. Unless otherwise noted, all preprocessing —including normalization, log-transform, and filtering —was performed using Scanpy^31^. Key datasets, platforms, and parameters are summarized below.

### Mouse small intestine dataset

The mouse small intestine spatial transcriptomics dataset was obtained from 10x Genomics’ public repository. It comprises a formalin-fixed paraffin-embedded (FFPE) mouse small intestine tissue section profiled with the Visium HD technology (high density 16 µm spots). For preprocessing, raw UMI counts were library-size normalized and log1p-transformed using Scanpy. Spots with <200 detected genes and genes present in <10 spots were removed.

The gene co-expression network was constructed using SpacGPA. Gene pairs with partial correlation (*Pcor*) ≥ 0.02 and co-expression in ≥ 10 spots were considered significantly co-expressed, yielding 53,566 significant gene pairs. MCL-Hub clustering (inflation=2, min module size=10) identified 61 gene programs. Program expression score were computed using the top 30 connectivity-ranked genes per program. GMM thresholding (posterior >0.9999, requiring ≥10 active spots per program) labeled active spots for each program, which were optionally smoothed as above. Spots were then assigned unique labels by expression ranking, with the optional local consensus step omitted. For validation, these programs were projected onto two mouse intestinal scRNA-seq datasets (Zwick et al. 2024^40^ and Xu et al. 2019^37^) by computing the same program intensity scores in the single-cell data.

### MOSTA dataset

The MOSTA dataset (Chen et al., 2022^10^) is a high-resolution spatial transcriptomic atlas of an entire E16.5 mouse embryo, denoted E16.5_E1S1, comprising 121,767 spots and 28,204 genes, covering all major organ regions of the developing embryo. Preprocessing followed the same procedure as above (normalization, log-transform, filtering spots with low gene counts and genes present in <10 spots). SpacGPA network inference, using a permutation-based FDR-selected *Pcor* threshold, retained 98,573 gene pairs, and MCL-Hub clustering identified 106 programs. Spatial expression profiles were then computed for all 106 programs in this section.

### Mouse pup dataset

The Mouse pup dataset is an in situ sequencing-based spatial transcriptomics dataset of an entire newborn mouse sagittal section. It was generated using 10x Genomics’ Xenium Prime 5K platform, profiling a targeted panel of ∼5,000 genes at single-cell resolution (∼1,300,000 cells in this sample). The gene-by-cell count matrix was loaded and log1p-transformed (library-size normalization was not applied). Cells with <100 detected genes were filtered out; all 5,010 genes were expressed in ≥10 cells, so no gene filtering was required. After filtering, 917,936 cells remained. SpacGPA network inference identified 37,318 significant gene pairs, and MCL-Hub yielded 88 programs. Program expression score were then computed for each cell.

### Mouse brain slices (Multi sections joint analysis)

Six adult mouse brain Visium sections (denoted Multi_train1–Multi_train6, from 10x Genomics) were analyzed jointly. Each section was normalized, log-transformed, and spot-filtered (>200 genes). We took the union of genes across all slices (22,277 genes expressed in ≥10 spots total) and concatenated the 16,603 spots (after QC) into a single matrix. Network inference with a partial-correlation cutoff (*Pcor*>0.032 after FDR control) yielded 59,919 gene pairs, from which MCL-Hub clustering detected 109 programs. These programs were then projected back to each individual section, as well as to three independent validation datasets: a high-resolution Visium HD brain slice, a Xenium Prime 5K brain slice, and a SMART-Seq single-cell brain dataset^64^.

### NSCLC dataset

The NSCLC dataset is a spatial transcriptomics dataset (Xenium Prime 5K platform) of a human non-small cell lung cancer (NSCLC) sample, specifically a lung adenocarcinoma section. It contains approximately 278,000 cells with spatial coordinates, covering both tumor epithelial cells and various tumor microenvironment (TME) cell types. After standard preprocessing (log-transforming counts and filtering low-quality cells), we constructed the gene co-expression network and applied MCL-Hub, identifying 58 programs, including ∼20 tumor-enriched programs. Program expression was computed for each cell, and cells were clustered in program-score space using (resolution 2.0), yielding 45 clusters.

Malignant clusters were defined based on multiple criteria: elevated expression of LUAD markers, high scores for cancer-related programs (cell cycle/proliferation/oncogenic pathways), spatial context (tumor core/invasive edge), and copy-number profiles inferred with infercnvpy^88^. Non-malignant cells with close spatial proximity to malignant cells were classified as TME. Tertiary lymphoid structures (TLS) were identified by gene-set scoring sing TLS signature, comprising germinal center B-cell genes (MS4A1, CD79A, CD79B); Tfh–B cell chemokine-axis genes (CXCL13, CXCR5, ICOS); dendritic-cell/high-endothelial venule markers (LAMP3, CD83); T-zone chemokines and receptors (CCR7, CCL19); and lymphoid-inducer genes (LTA, LTB)^89-93^. Top-scoring candidate spots were filtered by spatial connectivity, retaining contiguous regions above a minimum size. This two-stage criterion (molecular score plus spatial contiguity) yields TLS calls consistent with histological expectations.

### Comparison of SpacGPA with Other Methods

To evaluate SpacGPA’s performance in identifying gene programs, we compared it with five other methods on the same dataset (mouse small intestine). The methods included both matrix factorization and network/clustering approaches: NMF,cNMF, LDA, STAMP, and hdWGCNA^43-47^. For each method, we followed the recommended workflow or reference implementation. Where applicable, we set the number of programs/topics to match SpacGPA’s output. For NMF, LDA, and STAMP, we set the number of factors or topics to 61, equal to the number of programs SpacGPA found in this dataset. For cNMF, which estimates the optimal number of factors (K) internally, we used its built-in heuristic(yielding 40 programs). For hdWGCNA, which does not require pre-specifying the number of modules, we carefully tuned its parameters (e.g, network power, minimum module size, merging threshold) to obtain the best results; hdWGCNA ultimately detected 13 modules. The resulting programs from each method were evaluated using multiple criteria.

### Explained variance (expression reconstruction) evaluation

To assess how well each method’s programs capture data variability, we measured explained variance (EV). Let *X* be the *N* × *G* expression matrix (with *N* spots and *G* genes) and *M* an *N* × *K* matrix (with *N* spots and *K* programs). EV was evaluated as follows:

First, mean-center *X* by subtracting from each gene’s overall mean:

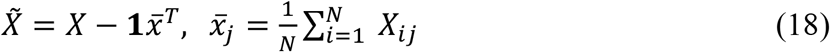

Next, solve for the coefficient matrix 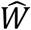 best reconstructs *X*′ from *M* in a least-squares sense:

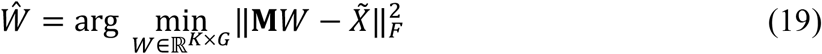

where | ⋅ |_*F*_ denotes the Frobenius norm. Using 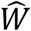, the reconstructed expression matrix is:

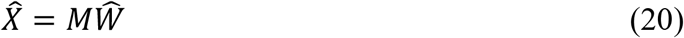

Then, define the residual sum of squares (*SS*_*res*_) and total sum of squares (*SS*_*tot*_) as:

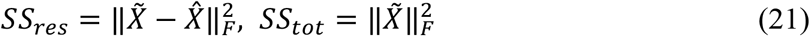

Finally, compute explained variance (EV) as:

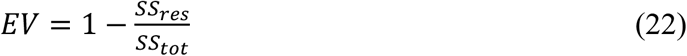

EV ranges from 0 to 1, indicating the fraction of total variance explained by the programs. Higher EV values mean the programs better capture the underlying expression structure.

### Spatial specificity evaluation

We compared the spatial clustering tendency of programs from each method by measuring their Moran’s I values. For each gene program, we computed Moran’s I on its spatial expression pattern (as described above in the *Spatial autocorrelation* section). Programs with higher Moran’s I exhibit stronger spatial localization, reflecting regions of either clustered high expression or clustered low expression.

### Biological interpretability evaluation

We assessed biological interpretability by performing GO enrichment on programs from each method using a unified pipeline. Two metrics were compared: (i) the average number of significant GO terms per program and (ii) the average number of non-duplicated GO terms per program (information diversity). To standardize gene set size, we used the top 30 genes per program ranked by each method’s weight matrix; for hard-clustering methods (SpacGPA, hdWGCNA) with <30 genes in a program, all genes were used. Metrics were computed as total significant (or deduplicated) GO terms across programs divided by the number of programs for that method.

### Visualization

We used several tailored visualization strategies. Co-expression networks were displayed in Cytoscape by exporting each program’s subnetwork and annotating nodes (e.g. highlighting hub genes or known markers)^94^. For functional enrichment, we generated bar plots of top GO/MP terms for each program (using SpacGPA’s module enrichment plotting), showing term significance and the contributing high-connectivity genes. Program–program similarities were visualized as heatmaps (via SpacGPA.module_similarity_plot) with hierarchical clustering based on Pearson and Jaccard distances, grouping programs with similar spatial profiles. We also plotted scatterplots of gene connectivity versus Moran’s I within each program to assess whether hub genes exhibit strong spatial patterning (via SpacGPA.module_degree_vs_moran_plot). Finally, program and gene expression were displayed on tissue coordinates (using scanpy.pl.spatial), and used dot plots (SpacGPA.module_dot_plot) were used to summarize expression across annotated cell types or regions.

### Quality Control

When constructing gene co-expression networks using SpacGPA, we used a permutation-based test to estimate the FDR of selected gene pairs. For Go or MP enrichment analyses, P values were computed using a one-tailed hypergeometric test and adjusted with the Benjamini-Hochberg method. In correlation analyses of connectivity and Moran’s I, multiple test was similary corrected using the Benjamini-Hochberg procedure. Ordinary least squares regression was applied to model connectivity as a function of Moran’s I, with 95% confidence intervals derived from the standard errors of the regression coefficients to quantify uncertainty in the fitted curve. When calculating the TLS marker gene module score with score_genes function in Scanpy, the function first bins all genes by their overall expression levels, then randomly selects control genes from each bin to construct a background set. The module score is defined as the mean expression difference between marker genes and background genes per cell. Finally, scores are Z-score normalized to reduce gene expression bias and improve the robustness of enrichment detechtion. All quantification and statistical analyses were performed in Python.

## Data and Code Availability

### Data availability

This study did not generate new experimental data. All datasets supporting the main findings are publicly available through existing databases or repositories. The specific datasets used (including their sources and accession information) are listed in the **Table S6**.

### Code availability

All data analysis was conducted on a computational workstation with 128 GB of memory and two Nvidia GEFORCE RTX 2080 Ti GPUs, as well as on a high-memory cluster node (2 TB RAM) for certain comparison methods. The SpacGPA toolkit, with documentation and usage examples, is freely available at https://github.com/MaShisongLab/SpacGPA. The complete code used to reproduce the analyses and figures in this study has been deposited in an online repository (to be released upon publication).

## Acknowledgements

We thank the USTC Supercomputing Center and USTC School of Life Sciences Bioinformatics Center for providing the computing resources.

## Author Contributions

**Yupu Xu:** Methodology, Investigation, Formal analysis, Writing – Original draft. **Lirui Chen:** Formal analysis. **Shisong Ma:** Conceptualization, Methodology, Formal analysis, Supervision, Writing – Original draft, Review & Editing, Funding acquisition.

## Supplemental Information

**Figure S1: Spatial expression and degree–Moran’s I concordance across 61 programs identified from the mouse small intestine dataset. (A)** Spatial expression of all 61 programs. **(B)** Scatter plots showing the relationship between degree and Moran’s I of genes within each program.

**Figure S2: Annotation integration of the mouse small intestine dataset using different methods**.

**Figure S3: Integrated annotation of the mouse small intestine dataset at 4 µm resolution using all programs except shared activity programs**.

**Figure S4: Spatial expression and degree–Moran’s I concordance across 106 programs identified from MOSTA dataset. (A)** Spatial expression of all 106 programs. **(B)** Scatter plots showing the relationship between degree and Moran’s I of genes within each program.

**Figure S5: Spatial expression across 88 programs identified from mouse pup dataset**.

**Figure S6: Spatial expression across 109 programs identified from joint analysis of six mouse brain coronal sections**. Expression of programs M1-M10 in all six sections **(A)** and other programs in the Multi_train1 section **(B)** are shown due to space limitations. Programs M12 and M98 are excluded in **(B)** because no more than 10 genes of these two programs were detected in Multi_train1 section.

**Figure S7: Spatial expression of all 109 programs from the joint analysis in an independent Visium HD dataset**.

**Figure S8: Spatial expression across 58 programs identified from a human lung adenocarcinoma sample**.

**Table S1: Summary of gene programs identified from the mouse small intestine dataset**.

**Table S2: Summary of gene programs identified from the MOSTA dataset**.

**Table S3: Summary of gene programs identified from the mouse pup dataset**.

**Table S4: Summary of gene programs identified from the joint analysis of six mouse brain coronal sections**.

**Table S5: Summary of gene programs identified from the human lung adenocarcinoma sample**.

**Table S6: Sources and accession numbers of public datasets used in this study**.

