## Supplemental Figures for "SpacGPA: annotating spatial transcriptomes through *de novo* interpretable gene programs"

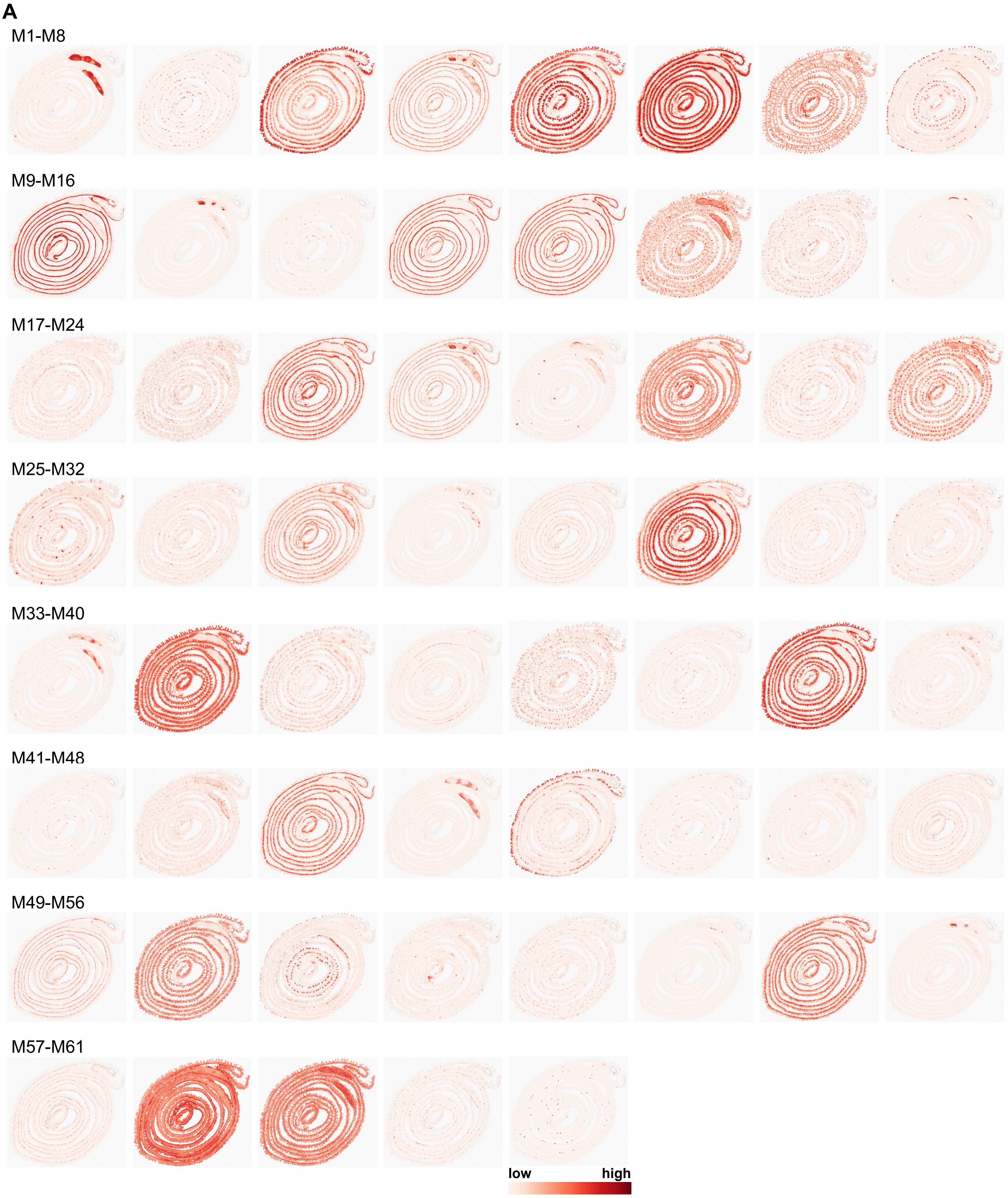
A
M1-M8
M9-M16
M17-M24
M25-M32
M33-M40
M41-M48
M49-M56
M57-M61
low
high

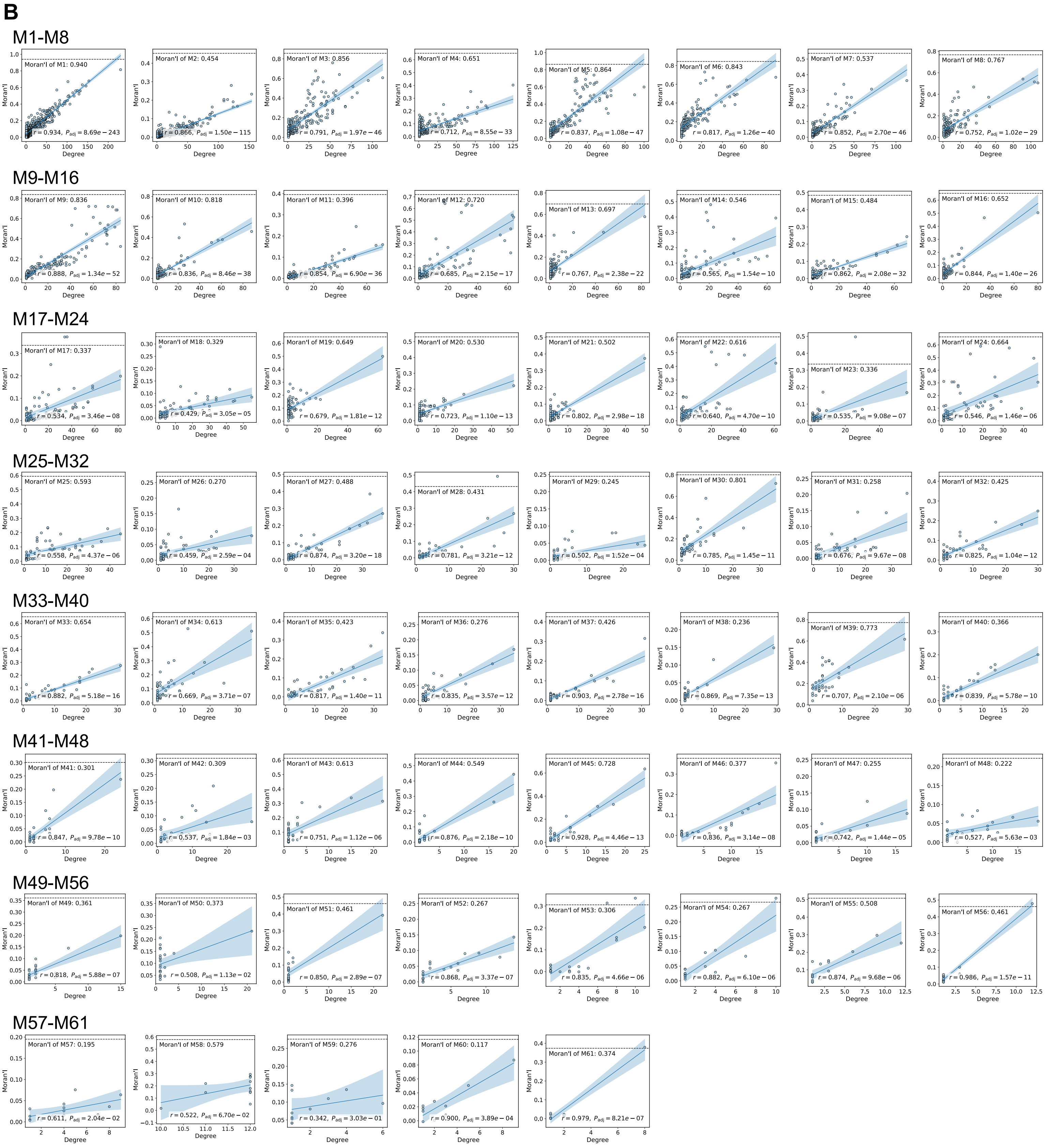


**Figure S1: Spatial expression and degree–Moran's I concordance across 61 programs identified from the mouse small intestine dataset. (A)** Spatial expression of all 61 programs. **(B)** Scatter plots showing the relationship between degree and Moran's I of genes within each program.

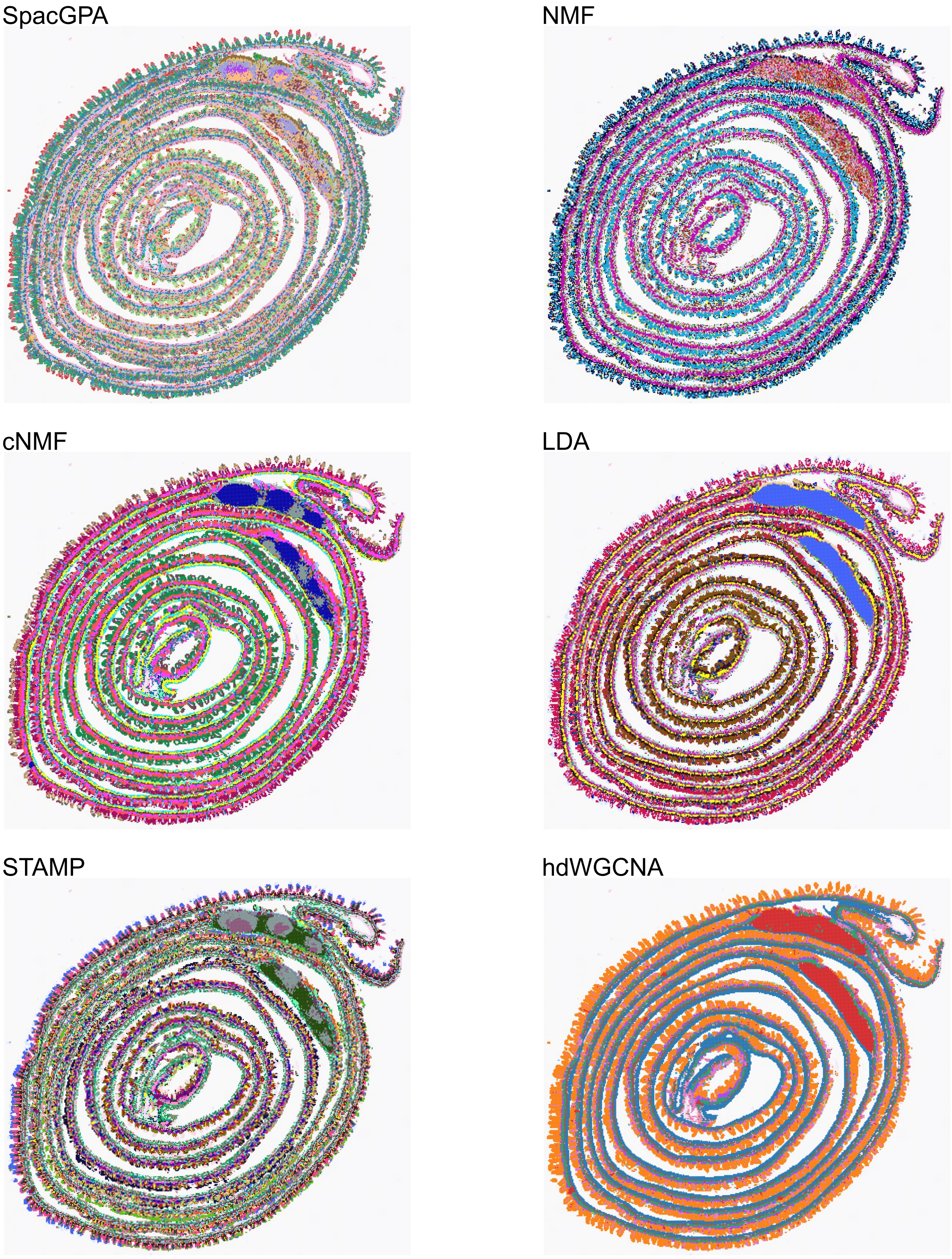


**Figure S2: Annotation integration of the mouse small intestine dataset using different methods.**

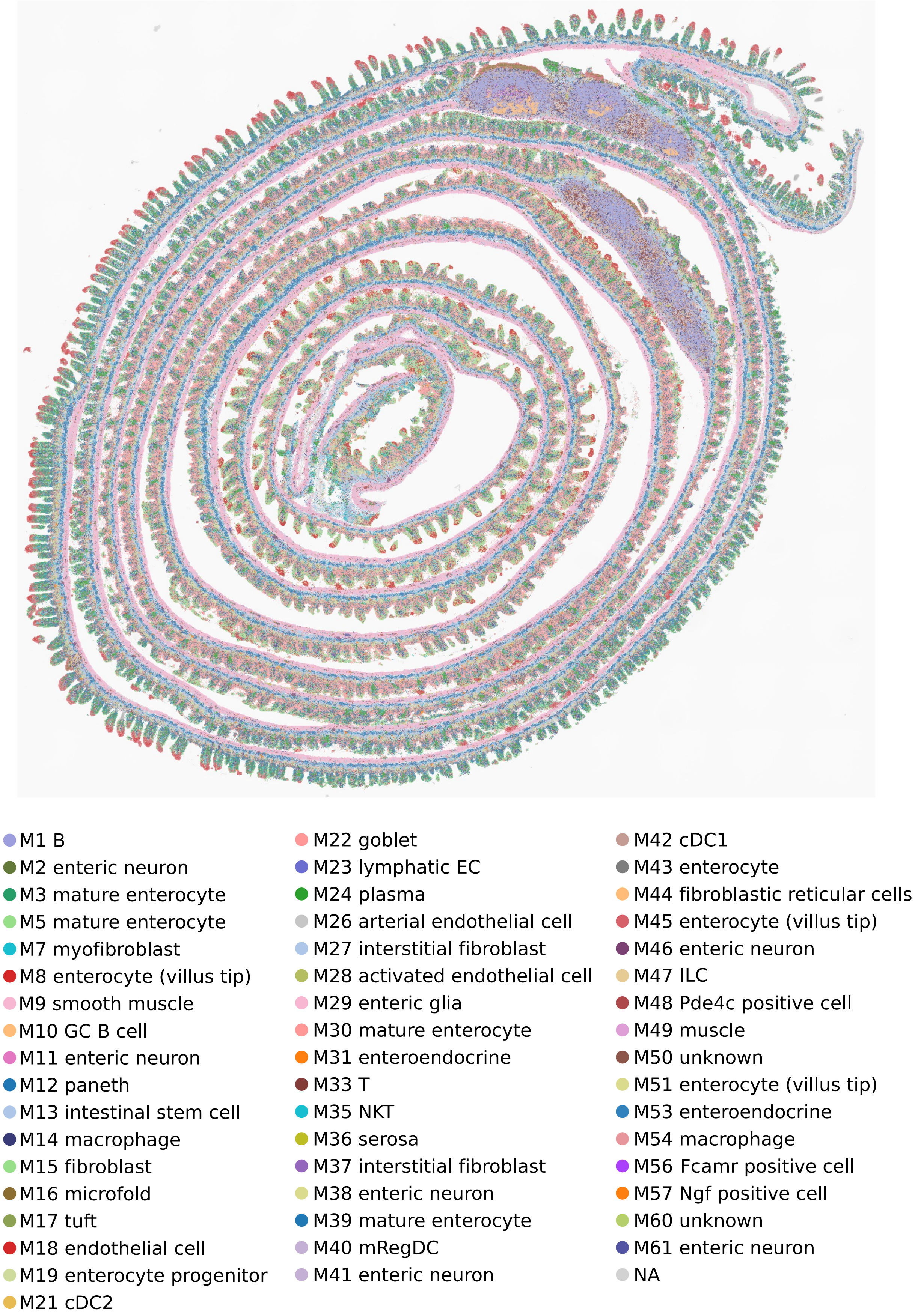


**Figure S3: Integrated annotation of the mouse small intestine dataset at 4 μm resolution using all programs except shared activity programs .**

A

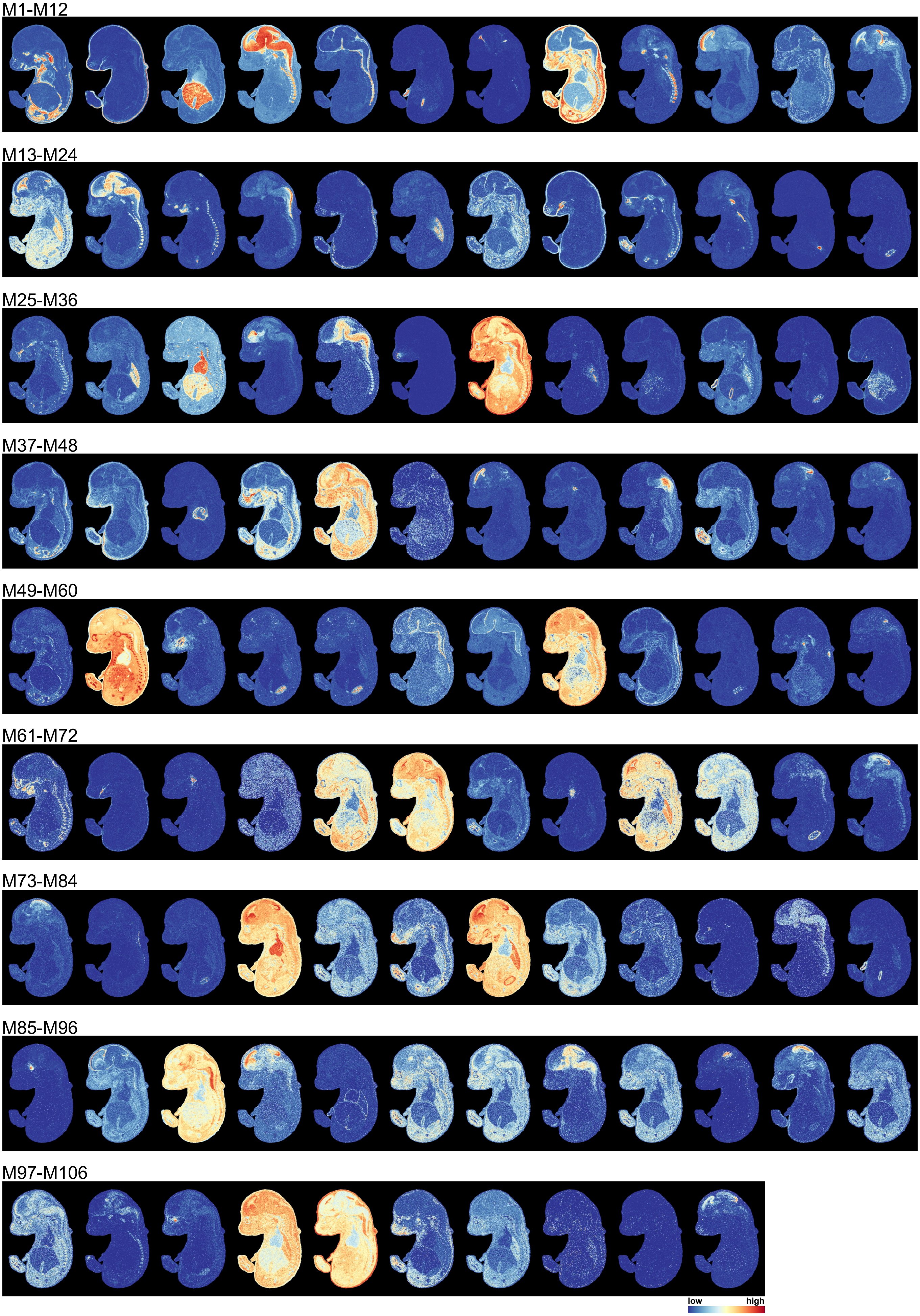

M1-M12
M13-M24
M25-M36
M37-M48
M49-M60
M61-M72
M73-M84
M85-M96
M97-M106
low
high

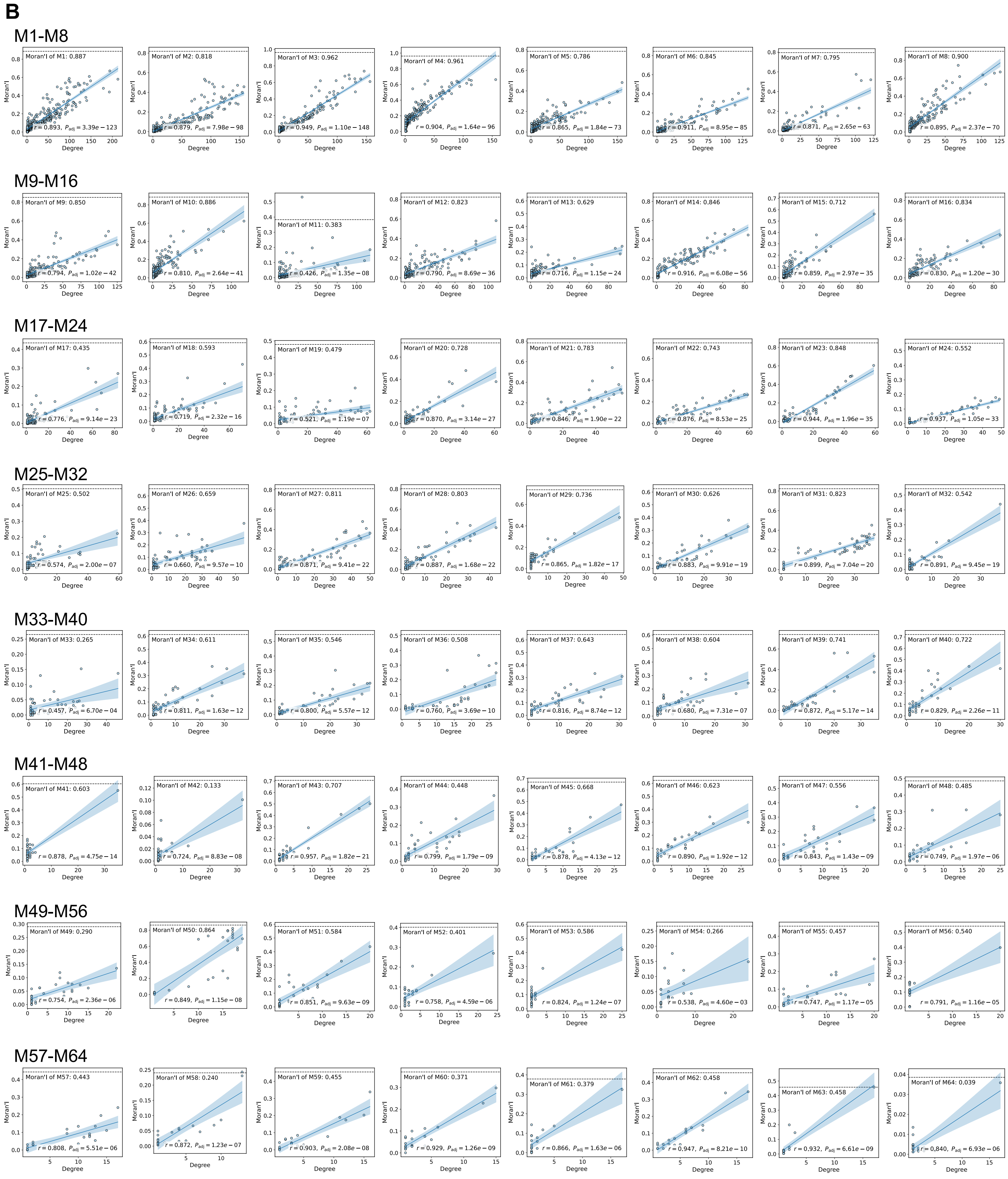
B
M1-M8
Moran'I of M1: 0.887
$r = 0.893$, $P_{adj} = 3.39e-123$
Moran'I of M2: 0.818
$r = 0.879$, $P_{adj} = 7.98e-98$
Moran'I of M3: 0.962
$r = 0.949$, $P_{adj} = 1.10e-148$
Moran'I of M4: 0.961
$r = 0.904$, $P_{adj} = 1.64e-96$
Moran'I of M5: 0.786
$r = 0.865$, $P_{adj} = 1.84e-73$
Moran'I of M6: 0.845
$r = 0.911$, $P_{adj} = 8.95e-85$
Moran'I of M7: 0.795
$r = 0.871$, $P_{adj} = 2.65e-63$
Moran'I of M8: 0.900
$r = 0.895$, $P_{adj} = 2.37e-70$
M9-M16
Moran'I of M9: 0.850
$r = 0.794$, $P_{adj} = 1.02e-42$
Moran'I of M10: 0.886
$r = 0.810$, $P_{adj} = 2.64e-41$
Moran'I of M11: 0.383
$r = 0.426$, $P_{adj} = 1.35e-08$
Moran'I of M12: 0.823
$r = 0.790$, $P_{adj} = 8.69e-36$
Moran'I of M13: 0.629
$r = 0.716$, $P_{adj} = 1.15e-24$
Moran'I of M14: 0.846
$r = 0.916$, $P_{adj} = 6.08e-56$
Moran'I of M15: 0.712
$r = 0.859$, $P_{adj} = 2.97e-35$
Moran'I of M16: 0.834
$r = 0.830$, $P_{adj} = 1.20e-30$
M17-M24
Moran'I of M17: 0.435
$r = 0.776$, $P_{adj} = 9.14e-23$
Moran'I of M18: 0.593
$r = 0.719$, $P_{adj} = 2.32e-16$
Moran'I of M19: 0.479
$r = 0.521$, $P_{adj} = 1.19e-07$
Moran'I of M20: 0.728
$r = 0.870$, $P_{adj} = 3.14e-27$
Moran'I of M21: 0.783
$r = 0.846$, $P_{adj} = 1.90e-22$
Moran'I of M22: 0.743
$r = 0.876$, $P_{adj} = 8.53e-25$
Moran'I of M23: 0.848
$r = 0.944$, $P_{adj} = 1.96e-35$
Moran'I of M24: 0.552
$r = 0.937$, $P_{adj} = 1.05e-33$
M25-M32
Moran'I of M25: 0.502
$r = 0.574$, $P_{adj} = 2.00e-07$
Moran'I of M26: 0.659
$r = 0.660$, $P_{adj} = 9.57e-10$
Moran'I of M27: 0.811
$r = 0.871$, $P_{adj} = 9.41e-22$
Moran'I of M28: 0.803
$r = 0.887$, $P_{adj} = 1.68e-22$
Moran'I of M29: 0.736
$r = 0.865$, $P_{adj} = 1.82e-17$
Moran'I of M30: 0.626
$r = 0.883$, $P_{adj} = 9.91e-19$
Moran'I of M31: 0.823
$r = 0.899$, $P_{adj} = 7.04e-20$
Moran'I of M32: 0.542
$r = 0.891$, $P_{adj} = 9.45e-19$
M33-M40
Moran'I of M33: 0.265
$r = 0.457$, $P_{adj} = 6.70e-04$
Moran'I of M34: 0.611
$r = 0.811$, $P_{adj} = 1.63e-12$
Moran'I of M35: 0.546
$r = 0.800$, $P_{adj} = 5.57e-12$
Moran'I of M36: 0.508
$r = 0.760$, $P_{adj} = 3.69e-10$
Moran'I of M37: 0.643
$r = 0.816$, $P_{adj} = 8.74e-12$
Moran'I of M38: 0.604
$r = 0.680$, $P_{adj} = 7.31e-07$
Moran'I of M39: 0.741
$r = 0.872$, $P_{adj} = 5.17e-14$
Moran'I of M40: 0.722
$r = 0.829$, $P_{adj} = 2.26e-11$
M41-M48
Moran'I of M41: 0.603
$r = 0.878$, $P_{adj} = 4.75e-14$
Moran'I of M42: 0.133
$r = 0.724$, $P_{adj} = 8.83e-08$
Moran'I of M43: 0.707
$r = 0.957$, $P_{adj} = 1.82e-21$
Moran'I of M44: 0.448
$r = 0.799$, $P_{adj} = 1.79e-09$
Moran'I of M45: 0.668
$r = 0.878$, $P_{adj} = 4.13e-12$
Moran'I of M46: 0.623
$r = 0.890$, $P_{adj} = 1.92e-12$
Moran'I of M47: 0.556
$r = 0.843$, $P_{adj} = 1.43e-09$
Moran'I of M48: 0.485
$r = 0.749$, $P_{adj} = 1.97e-06$
M49-M56
Moran'I of M49: 0.290
$r = 0.754$, $P_{adj} = 2.36e-06$
Moran'I of M50: 0.864
$r = 0.849$, $P_{adj} = 1.15e-08$
Moran'I of M51: 0.584
$r = 0.851$, $P_{adj} = 9.63e-09$
Moran'I of M52: 0.401
$r = 0.758$, $P_{adj} = 4.59e-06$
Moran'I of M53: 0.586
$r = 0.824$, $P_{adj} = 1.24e-07$
Moran'I of M54: 0.266
$r = 0.538$, $P_{adj} = 4.60e-03$
Moran'I of M55: 0.457
$r = 0.747$, $P_{adj} = 1.17e-05$
Moran'I of M56: 0.540
$r = 0.791$, $P_{adj} = 1.16e-05$
M57-M64
Moran'I of M57: 0.443
$r = 0.808$, $P_{adj} = 5.51e-06$
Moran'I of M58: 0.240
$r = 0.872$, $P_{adj} = 1.23e-07$
Moran'I of M59: 0.455
$r = 0.903$, $P_{adj} = 2.08e-08$
Moran'I of M60: 0.371
$r = 0.929$, $P_{adj} = 1.26e-09$
Moran'I of M61: 0.379
$r = 0.866$, $P_{adj} = 1.63e-06$
Moran'I of M62: 0.458
$r = 0.947$, $P_{adj} = 8.21e-10$
Moran'I of M63: 0.458
$r = 0.932$, $P_{adj} = 6.61e-09$
Moran'I of M64: 0.039
$r = 0.840$, $P_{adj} = 6.93e-06$
Moran'I
Degree

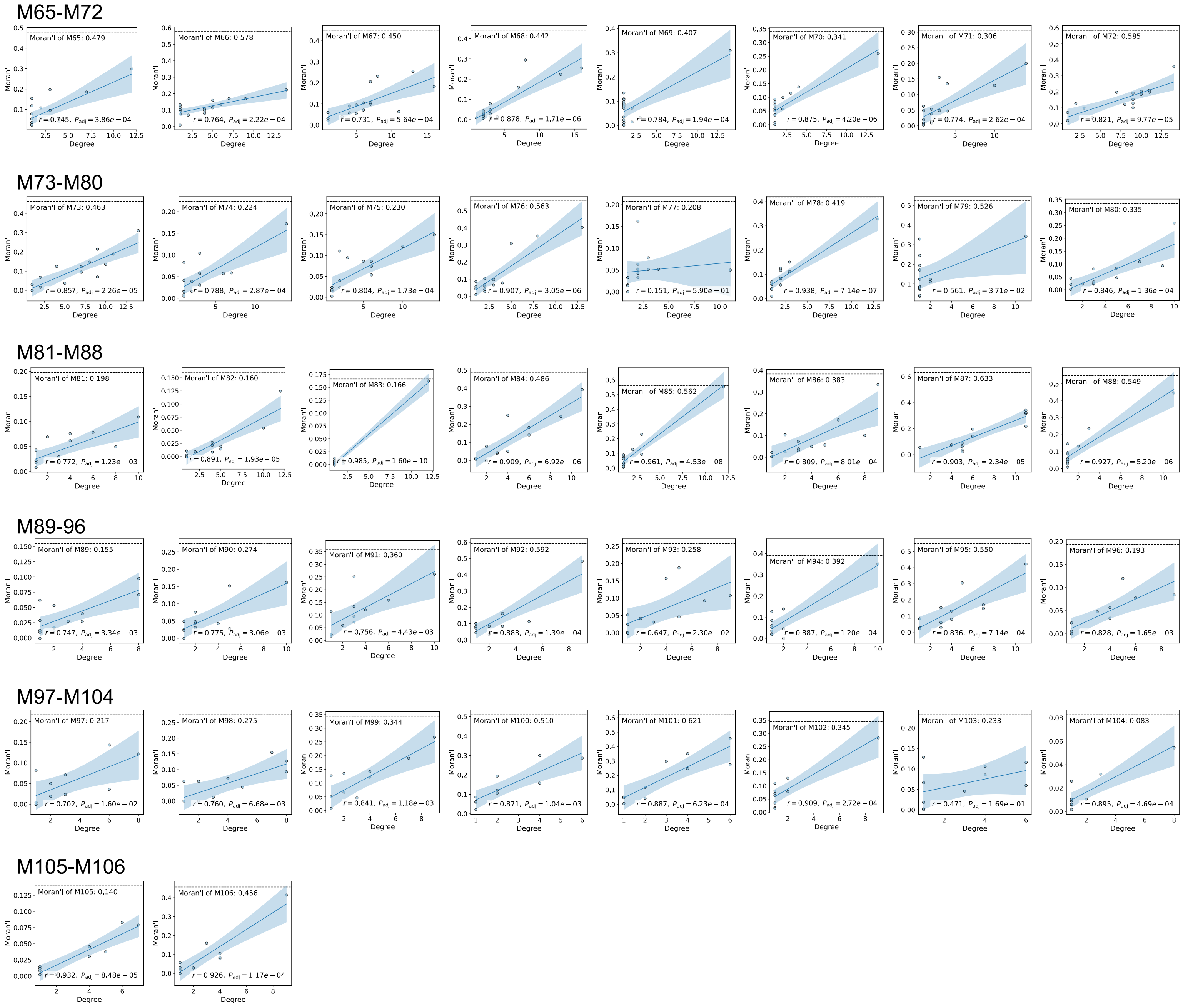


**Figure S4: Spatial expression and degree–Moran's I concordance across 106 programs identified from MOSTA dataset. (A)** Spatial expression of all 106 programs. **(B)** Scatter plots showing the relationship between degree and Moran's I of genes within each program.

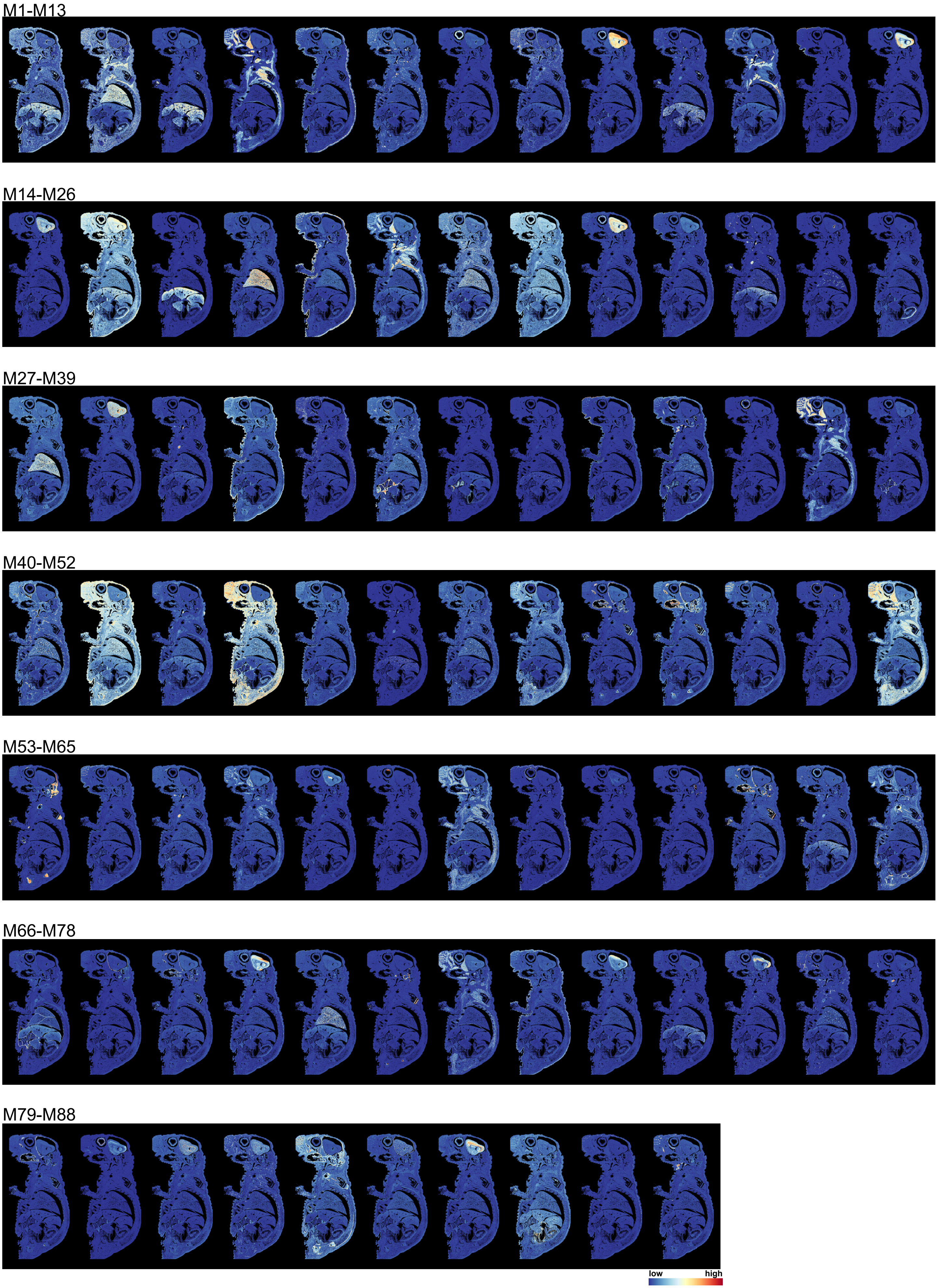


**Figure S5: Spatial expression across 88 programs identified from mouse pup dataset.**

**A**

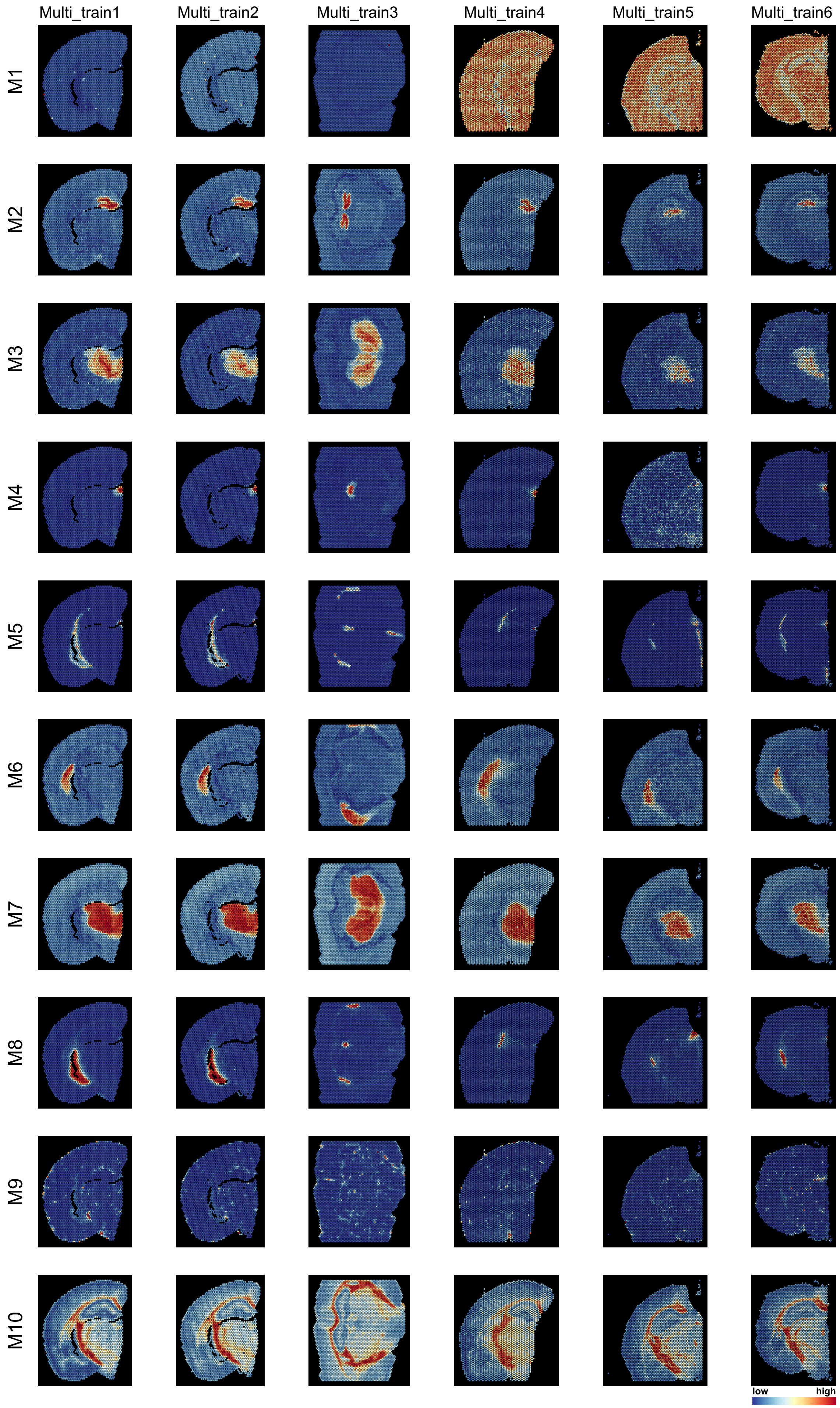

Multi_train1
Multi_train2
Multi_train3
Multi_train4
Multi_train5
Multi_train6
M1
M2
M3
M4
M5
M6
M7
M8
M9
M10
low
high

B

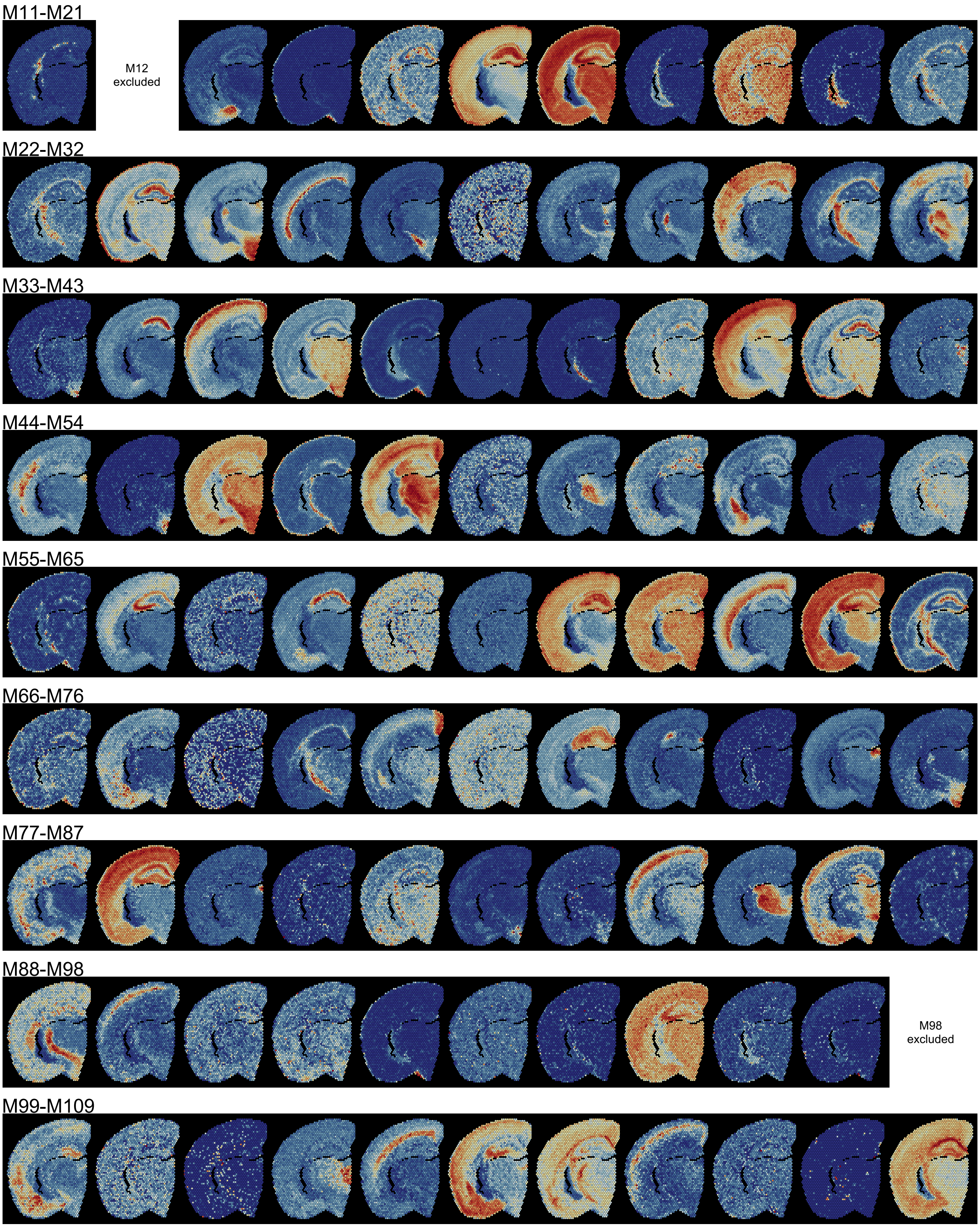


**Figure S6: Spatial expression across 109 programs identified from joint analysis of six mouse brain coronal sections.** Expression of programs M1-M10 in all six sections **(A)** and other programs in the Multi_train1 section **(B)** are shown due to space limitations. Programs M12 and M98 are excluded in **(B)** because no more than 10 genes of these two programs were detected in Multi_train1 section.

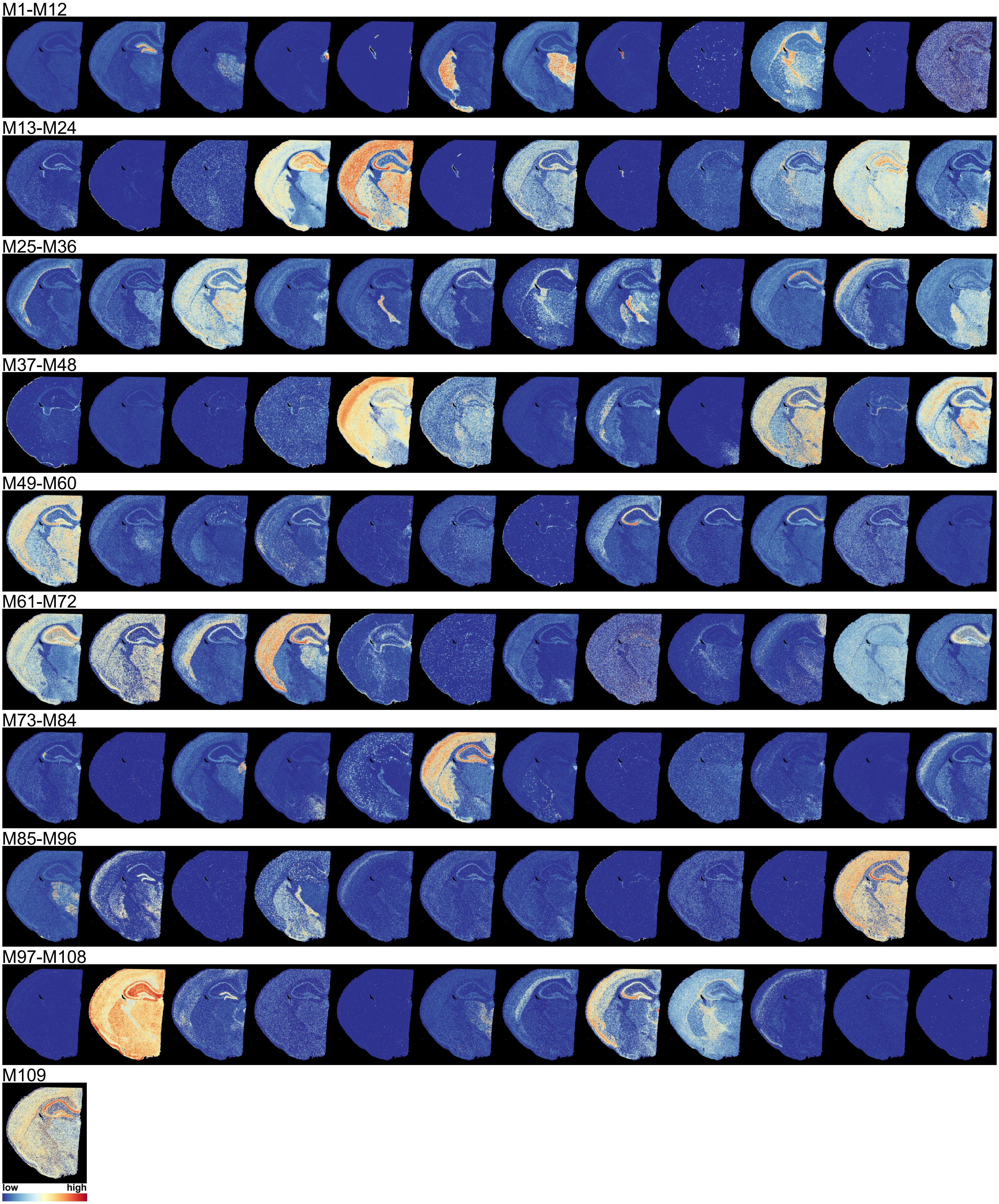


**Figure S7: Spatial expression of all 109 programs from the joint analysis in an independent Visium HD dataset.**

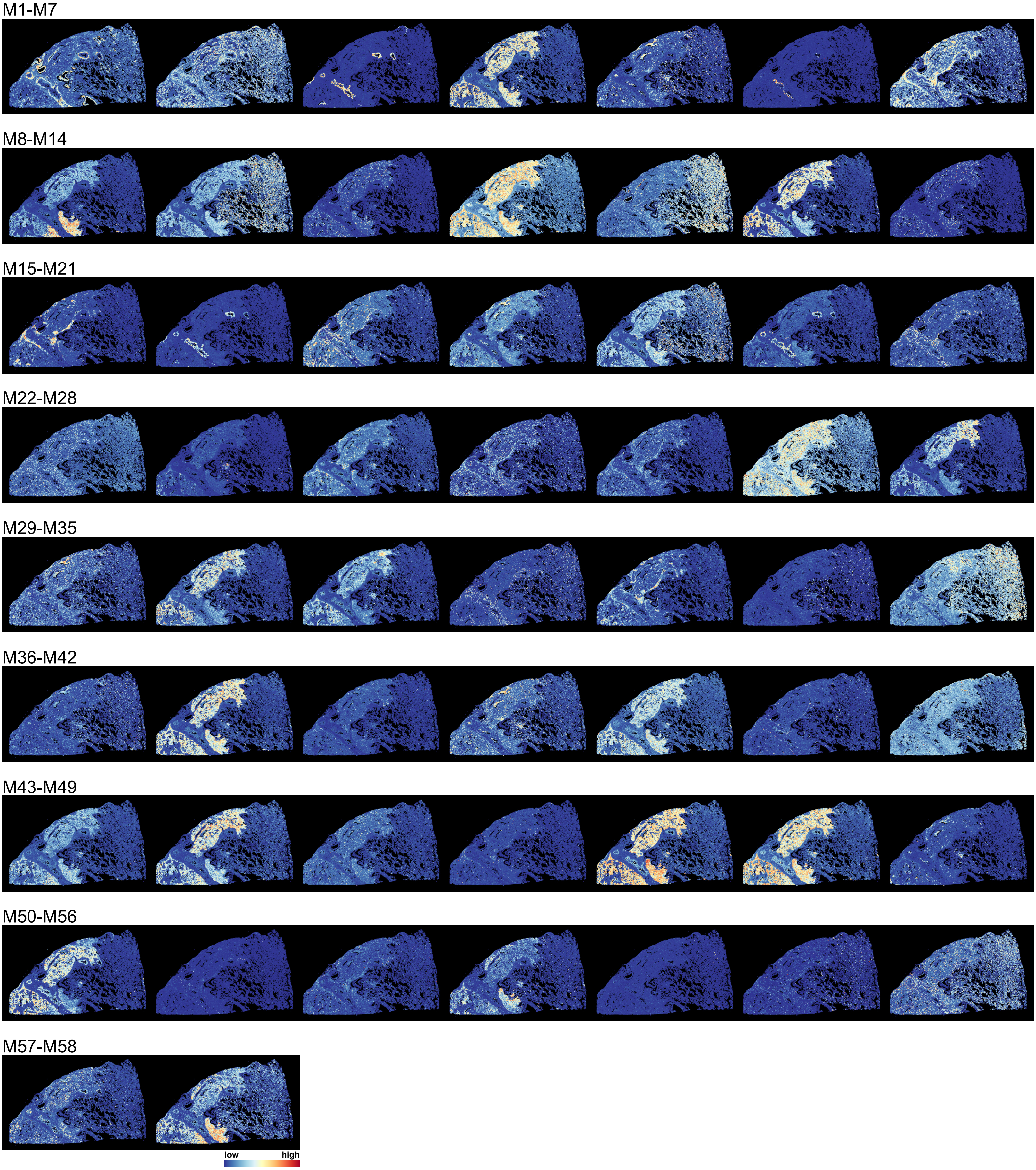


**Figure S8: Spatial expression across 58 programs identified from a human lung adenocarcinoma sample.**
